# A Daily Cycle of White Collar Complex Dephosphorylation Sustains Circadian Rhythmicity in *Neurospora*

**DOI:** 10.1101/2025.09.25.678408

**Authors:** Bin Wang, Xiaoying Zhou, Jennifer. J. Loros, Jay C. Dunlap

## Abstract

As a photoreceptor, the transcription factor complex WCC acutely activates ∼5% of the genome in response to blue light, while as the circadian positive element in the dark WCC influences expression of about 40% of the transcriptome. Among WCC-regulated genes is *frq* which is both acutely light-activated through a *pLRE* and circadian-regulated through a *C-box* promoter element that is not active in constant light. The complex of FRQ, FRH, and CK1, the FFC, phosphorylates WCC at >95 sites, thereby repressing its activity and closing the circadian feedback loop in the dark. Although FFC has no described role in the light, we validated the expectation that FFC-driven WCC phosphorylation also silences *C-box* promoters in constant light, thereby confirming two classes of WCC targets, *C-box*-like that are normally repressed in the light and *pLRE*-like that remain light-active despite FFC-driven WCC phosphorylation. Genome-wide derepression of *C-box-*like promoters in *frq*-null fungi may explain reported non-circadian effects seen in some *frq*-null fungi including reduced virulence and conidiation. Reanalysis of WCC-mediated circadian activation and repression revealed that, while at dusk most WCC is phosphorylated and repressed, subsequent circadian activation is the result of transient dephosphorylation/derepression of just a small subset of this WCC pool; this small active pool drives expression of FRQ, nucleating the FFC, which rapidly re-phosphorylates the WCC pool to repress it, a phosphorylation/dephosphorylation cycle that can run for days without new WCC synthesis. The realization that both FFC and WCC are regulated primarily through phosphorylation rather than turnover leaves the circadian oscillator looking much like a phoscillator, emphasizing the primacy of post-translational regulation in timekeeping.

**Significance:** At the core of circadian clocks of fungi and animals, a protein heterodimer drives expression of gene(s) whose products inactivate the heterodimer via phosphorylation. To sustain the cycle through multiple days, the activity of the heterodimer must be restored, but the means through which this happens have been unclear. In the clock model *Neurospora*, the White Collar Complex (WCC) is the heterodimer and FFC is the complex that inactivates it. We determined that WCC activity is restored principally by removal of the inhibitory phosphorylations and that for at least several days no new synthesis of WCC is required. The results confirm the existence of a large pool of inactive WCC in the cell and highlights the delicate balance between FCC-dependent phosphorylation/inactivation and phosphatase-dependent dephosphorylation/reactivation. Each morning, this balance allows transient activation of a fraction of the inactive WCC pool, thereby restarting the circadian cycle.

## Introduction

Circadian clocks regulate a wide variety of behavioral, physiological, cellular, and molecular events (1). Dysfunction of biological rhythms has been linked to various human diseases (2, 3). The core oscillators in fungi and animals are intracellular, constructed as autoregulatory transcriptional-translational negative feedback loops wherein a heterodimer of positive elements drives expression of negative elements, and the latter repress transcriptional activity of the former to ultimately impinge on their own expression (4–6). For instance, in mammalian cells, transcription factors CLOCK and BMAL1 heterodimerize via PAS domains and drive expression of negative elements, PERs and CRYs, as well as elements in ancillary loops that contribute to robustness (5, 7). In *Neurospora* transcription factors White Collar 1 (WC-1) and WC-2 heterodimerize via PAS domains to form the White Collar Complex (WCC), which drives expression of the core negative regulator FRQ under contrasting light conditions: WCC either binds to the *proximal Light-Response Element* (*pLRE*) to mediate *frq*-induction in the light (8, 9), or binds rhythmically to the *Clock-box* (*C-box*) in the dark, driving the cyclical expression of *frq* required for rhythmicity (10–13). FRQ interacts with FRH (FRQ-Interacting RNA Helicase) (14–16) and CK1 (casein kinase 1) (12, 17, 18) to form the FFC (FRQ and FRH complex), which in turn represses circadian (but not light-driven) activity of WCC to terminate its own expression, thereby closing the circadian feedback loop (4, 7, 17).

Multisite phosphorylation has been found to be a central mechanism in tightly controlling and fine-tuning circadian functions of both the positive and negative elements in these loops (7). Regulation of the negative elements in animals and *Neurospora* has received the most attention as their time-of day phosphorylation (17, 19–22) leads directly to their loss of activity, although there is debate about whether phosphorylation first leads to inactivation followed by turnover as has been shown in *Neurospora* (23, 24), or if inactivation is caused by turnover elicited by phosphorylation as is believed for mammalian clocks (5). FRQ, for instance, undergoes myriad temporal phosphorylations by CK1, CK2 and additional kinases with over 100 time-of-day-specific phosphorylations, tightly tuning its activity, dynamically determining its binding partners, and eventually resulting in its inactivation (17, 21, 22) which happens prior to its turnover (23). Positive element heterodimers in both fungi and mammals are also subject to time-of-day specific phosphorylation leading to repression by reducing their ability to bind to DNA (25–31) (Displacement Repression (29)). In *Neurospora* > 80 phosphosites have been identified on WC-1 and 15 on WC-2 (28). Circadian repression requires phosphorylation of distinct clusters of these residues on each protein, and the time-of-day-specific phosphocode governing rhythmic repression has been elucidated (28). Very recent comparable data from mammalian cells has identified sites on both BMAL1 and CLOCK that must be phosphorylated to achieve repression (32), a result congruent with that seen in *Neurospora* thus establishing a conserved precedent for repression.

Aside from effects of kinases and phosphorylation, few studies have assessed dosage requirements for circadian positive elements or the role(s) of phosphatases in influencing this. In *Neurospora*, period length is little affected in strains constitutively or overexpressing WC-1, WC-2, or both together (14, 33), and likewise mammalian cells expressing various constitutive levels of BMAL1 or CLOCK are strongly rhythmic with periods similar to WT (31, 32). Although rhythms in positive element expression are seen both in *Neurospora* WCC (34–36) and in mammalian BMAL1 (e.g. (27)), the rhythms are not viewed as essential but instead as contributing to robustness of the cycle (e.g. (37)), in all suggesting that while the circadian heterodimer (WCC or BMAL1/CLOCK) is necessary for rhythmicity, neither its rhythmic expression nor abundance acts as a causal element in determining the pace of the core clock. This, however, poses a question: If the cycle of positive element activity required for the clock depends partly (as in mammals) or wholly (as in *Neurospora*) on a cycle of phosphorylation-mediated repression, how can the cycle continue unimpeded in the presence of a steady influx of new, unmodified and non-repressed heterodimer (e.g. (14))? Such an influx of active positive elements ought to short-circuit the feedback loop leading to constant negative element expression and loss of rhythmicity.

Phosphatases that can reverse the effects of critical clock kinases such as CK1 and CK2 have been suggested as likely clock-relevant proteins and could address this paradox in fungi and mammals, but the phenotypes of mutants are not as unambiguous as could be hoped. Downregulation of PP1 or PP2A lengthens period of the *Drosophila* clock (38, 39); in fibroblasts PP1 knockdown results in period shortening (40) whereas it is reported to lengthen period in U2OS cells (41), and in fibroblasts reduced PP5 caused modest period lengthening (42). In all cases effects were interpreted as affecting phosphorylation of Per proteins, not their heterodimeric activators. In U2OS cells, PPP4 has been shown to dephosphorylate BMAL1 but paradoxically, elevated PPP4 activity (which results in hypophosphorylation of BMAL1 and enhanced DNA-binding activity) results in reduced transactivation activity with a slightly longer period length, as it does in *Drosophila* (43). Conversely in *Neurospora*, the *rgb-1* mutant in the essential regulatory subunit of PP2A causes a reduction in PP2A activity leading to WCC hyperphosphorylation, reduced DNA binding causing less active *frq* expression and a three-hr period lengthening with no loss of rhythm strength (25, 28), whereas reduction/loss of PPP-1 or PPH-4 yields a 2-3 hr period shortening (44, 45). Thus in both mammalian and fungal cells the rhythm continues despite loss of salient phosphatase activities (28, 44) suggesting redundancy or a modulatory as distinct from an essential role for dephosphorylation; notably, however, even the phenotypes resulting from loss of phosphatases lack internal consistency. This difficulty in drawing simple conclusions from correlations between phosphatase activities, positive element heterodimer function, and period length is highlighted by the Δ*csp-6* strain in *Neurospora* that lacks the CSP-6 phosphatase and in which, while WCC is always hyperphosphorylated, the core clock still maintains a normal circadian period (46). Taken together, these inconsistencies suggested that there is more to find out about how positive element heterodimers in general, or WCC in particular, renews its activity to sustain circadian rhythms. We find, confirming prior predictions (25), that the answer lies in a more nuanced view of the pool(s) of WCC in the cell: At all times in both light and dark, most of the WCC pool is phosphorylated and inactive at *C-box* promoters; activation of negative element (here *frq*) expression relies on dephosphorylation/reactivation of only a small fraction of this cellular WCC pool, which becomes transiently dephosphorylated at times when WCC activity peaks in driving circadian expression of *frq*. We find the circadian cycle of WCC repression and reactivation does not require ongoing transcription or translation, and inhibition of protein translation in the cell does not block robust dephosphorylation/reactivation of WCC (28), and even when the induction of *wc-1*, *wc-2*, or both is turned off, existing WCC is still able to support sustained circadian oscillations. These data provide an enhanced mechanistic basis for understanding how phosphorylation/dephosphorylation cycles lead to cyclic repression and re-activation of a heterodimeric transcriptional complex driving a circadian oscillation, highlighting significance of the balance between phosphorylation and dephosphorylation of circadian positive elements and thereby illuminating molecular events lying at the core of the circadian oscillator.

## Results

### An activity for FRQ in the light

As a complex sensing light, WCC undergoes conformational changes to form a homodimer, strongly inducing FRQ expression mainly through the *pLRE* element in the *frq* promoter (9). In the dark, WCC acts simply as a heterodimer and FRQ complexes with FRH and CK1, the FFC, to repress the circadian transcriptional activity of WCC by promoting its phosphorylation (12, 15, 17, 25, 28). FRQ and FFC have described roles only in the circadian system in darkness where, by repressing WCC activity, they impact expression of ∼40% of the transcriptome (34); however, we find that FFC also represses WCC transcriptional activity at *C-box* in the light, an activity that is not surprising in hindsight.

Luciferase (Luc) expression driven by the *C-box* was tracked in *ffc* mutants Δ*frq* and *frh^R806H^*, an *frh* mutant deficient in WCC repression in the dark (14). WCC activity at the *C-box* in either Δ*frq* or *frh^R806H^* was prominently enhanced at the L-to-D transition (Fig. 1A), confirming that FFC also functions in the light to inhibit WCC at the *frq* promoter. Because WCC is known to bind to and regulate myriad genes during the circadian cycle in darkness that are normally repressed in light, the finding that FFC also represses this potentially light-driven promoter activity has biological significance: In strains lacking FRQ, *C-box*-regulated genes could be highly expressed in the light. To confirm that the novel activity of FRQ in the light on *C-box*-like promoter elements reflected the absence of FRQ/CK1-mediated phosphorylation/repression of WCC and did not require light-driven de novo synthesis of WC-1, we examined two direct WCC targets with *C-box*-like promoters, NCU02713 (*csp-1*) and NCU09841 (*pty-4)*, whose transcription is reduced in the absence of functional WCC (Fig. S1A and S1C). Utilizing a Δ*frq*, *qa-2:wc-1* strain (Fig. S1B) in which synthesis of WC-1 is dependent on QA and not light, we saw that once WC-1 was induced it led to sustained elevated target gene expression (Figs. S1C and S1D), consistent with the idea that, under light conditions, FFC represses WCC transcriptional activity at *ccg* promoters.

**Fig. 1.**
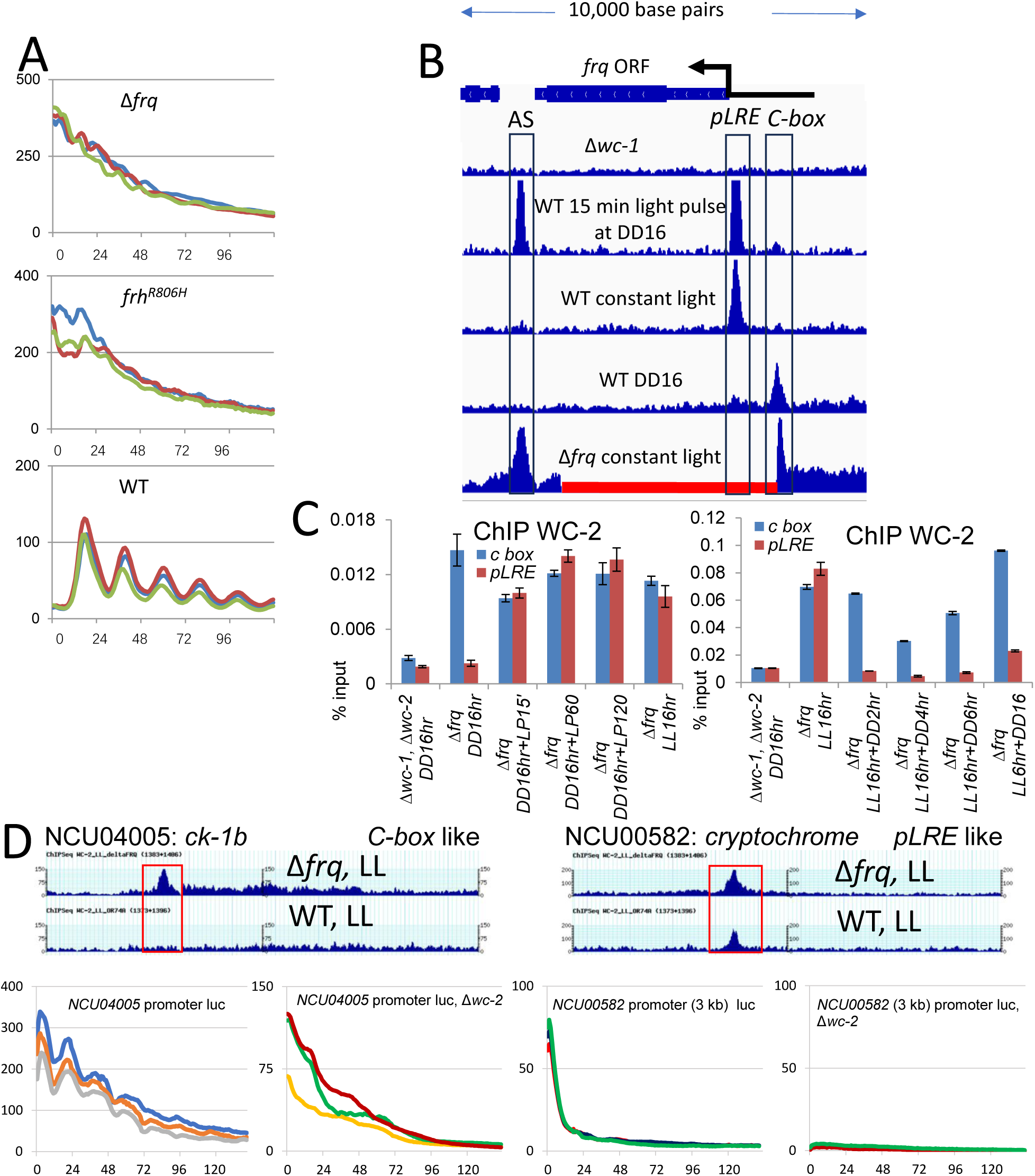
Enhanced *frq C-box* promoter activity of *frq* and *frh* mutants in the light reveals a function for FRQ in light-signaling. (**A**) *C-box* promoter-driven *luciferase* expression was tracked in real-time in Δ*frq* and *frh^R806H^*. Strains were grown in light overnight at 25 °C, synchronized by a light-to-dark (L-D) transfer, and bioluminescence signals were followed hourly in darkness at the same temperature. Three replicates (in three different colors) are shown with the x- and y-axes representing time (in hours) and signal intensity (in arbitrary units), respectively. (**B**) ChIP-seq using anti-WC-2 antibody as a proxy for WCC binding at the *frq* promoter in strains and conditions as marked. The gene map of the *frq* locus is at the top; the black left-facing arrow marks the transcription start site, the *C-box, pLRE,* and AS (antisense) promoter elements are marked, and the Δ*frq* deletion is marked with the red bar at the bottom; this deletion is the standard *frq* deletion from the *Neurospora* knockout collection (75). Published ChIP data for constant light and DD16 (35) are included for reference. (**C**) In Δ*frq, pLRE* retains light-induction that is lost in dark whereas *C-box* is always active. ChIP-quantitative PCR was carried out using WC-2 antibody with *C-box*- or *pLRE*-specific primer sets using strains and conditions as marked. Average values were plotted as a percentage of the total; error bars show the standard error of the mean (SEM [n = 3]). Signals obtained from ChIP were divided by input signals and are reported as percentage of inputs. Three technical replicates were performed, and the bars represent average values plotted as a percentage of the input, with error bars representing the SEM (n = 3). Note that these experiments employed a novel Δ*frq* strain engineered to remove only the ORF while leaving *C-box* and *pLRE* promoter sequences intact. (**D**) Examples of a *C-box-like* promoter (WCC-bound in Δ*frq* but not in WT in LL; *frq* and WCC-dependent rhythmicity in DD) and *pLRE*-like promoter (WCC binding not *frq-*responsive in LL; arrhythmic in DD). (Top) WC-2 ChIP-seq data covering the *C-box* in Δ*frq* (top) and WT (bottom) strains grown in constant light. NCU04005 (*casein kinase-1b* [*CK-1b*]) is an example of a *C-box*-like promoter while NCU00582 (*cryptochrome* [*cry*]) is an example of a *pLRE*-like promoter. (Bottom) Promoters of indicated genes were individually fused to the *luciferase* gene, the constructs were transformed to the *cyclosporin resistance*-*1* (*csr-1*) locus, and light signals from the transgenic strains were followed in the dark over five days. “Δ*wc-2*” means in the genetic background of *wc-2*-null. Y-axis marks light intensity in arbitrary units; x-axis refers to time in hrs.

To probe the effect of FRQ on WCC activity in the light, we compared binding of WCC to the *C-box*, *pLRE*, and *frq* antisense promoter in Δ*frq* upon light exposure in a chromatin-immunoprecipitation assay followed by sequencing (ChIP-seq). In WT following a brief light pulse given in darkness, WCC binds strongly to the *pLRE* and *frq* antisense promoters (Fig. 1B; see also (47)) whereas binding to the *C-box* is greatly reduced (Fig. 1B, (47)); in constant light, binding to the *frq* antisense promoter is lost but *pLRE* binding remains. However, in Δ*frq* grown in constant light, WCC binds strongly to *C-box* and *frq* antisense promoters despite the light (Fig. 1B), confirming that FFC removes WCC from the *C-box* in the light as it does in the dark. Because the *frq* deletion used in Fig. 1B removes the *pLRE*, RT-qPCR was used to track WCC binding to the *pLRE* and *C-box* in a different *frq* deletion strain that just removes only coding sequences while leaving both sense promoters intact (Fig. 1C). Loss of FRQ renders binding to *C-box* always elevated but does not impact light-regulation of *pLRE* binding: Light still induces *pLRE* binding (left panel, Fig. 1C) and darkness lessens *pLRE* binding (right panel, Fig. 1C). In constant light, WCC binds strongly to *pLRE* despite high abundance of FRQ in the cell (48, 49), whereas FRQ (presumably with FRH and CK1) represses association of WCC with the *C-box* and the antisense promoter. Taken at face value these data indicate that there are potentially two classes of WCC-responsive directly light-regulatable promoters in the genome, *pLRE*-like that have been previously described (e.g. (47, 48)) and *C-box*-like, repressed in the light and cyclically active in the dark. In ChIP-seq experiments we examined WCC binding genome-wide in WT and in Δ*frq* strains and identified both classes, exemplars of which are shown in Fig. 1D, with a more extensive list see in Fig. S1A and compiled in Suppl. Table 1. We find that roughly half of genomic sites bound by WCC fall into each class.

### Unphosphorylated WCC appears when its transcriptional activity peaks over a circadian cycle

A prediction of the data in Fig.1, especially as regards *frq*, is that in order to keep *C-box*-like promoters repressed at the start of night, essentially the entire pool of WCC must be phosphorylated in the light at the Ser/Thr residues found to be essential for circadian repression (28). We tested this hypothesis using Phos-tag gels to distinguish a small fraction of unphosphorylated WC-1 from the much larger expected pool of phosphorylated WCC (Fig. 2A). The phosphosites of WC-1 that are collectively indispensable for feedback loop closure are S971, S988, S990, S992, S994, and S995; in *wc-1^1-970A&996-1167A^* all identified WC-1 phosphosites except for S971, S988, S990, S992, S994, and S995 were collectively mutated to alanines; this strain still supports a robust oscillator ((28) and Fig. 2A, top). As predicted, at the time of the L-to-D transition, all the essential residues of WC-1 and most of WC-2 are phosphorylated such that the WCC is inactive with respect to *C-box* activity (Fig. 2A, middle, Phos-tag data). Following the L-D transfer, the amount of WC-1 appears relatively constant, but strikingly most of the cellular pool of WC-1 remains fully phosphorylated throughout the day (Fig. 2A, middle). A fraction of this pool becomes dephosphorylated apparently synchronously at all six sites beginning sharply at CT22 (DD8, 8 hrs into darkness), due either to new synthesis or dephosphorylation, peaking 4 hrs later and disappearing by CT14 (DD24) (Fig. 2A). These changes are congruent in timing and magnitude with the rhythmic binding of WCC to the *C-box* that drives *frq* transcription throughout a day (Fig. 2B; (35, 47)).

**Fig. 2.**
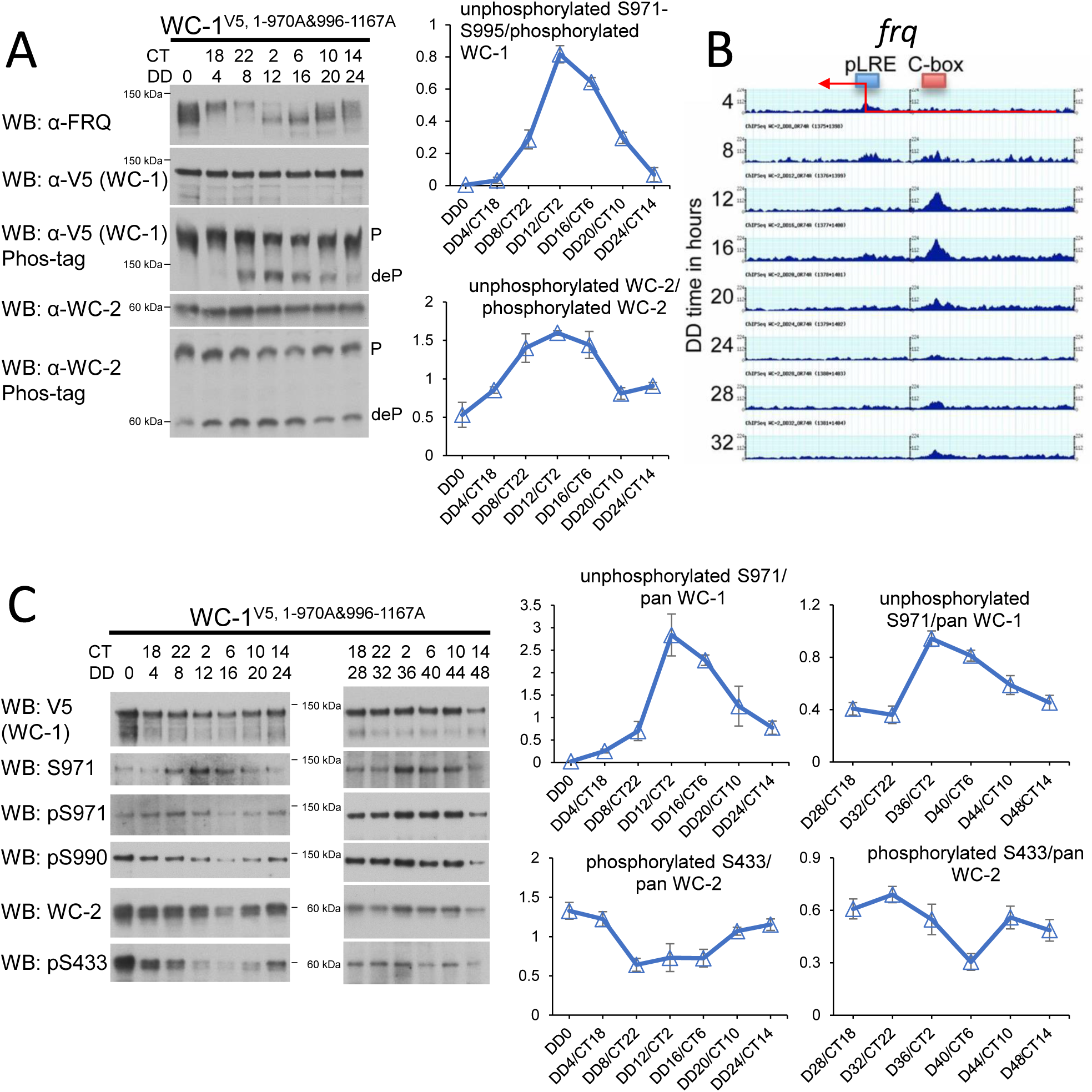
A small fraction of WC-1 and WC-2 become dephosphorylated coincident with FRQ expression. (**A**) Phosphorylation at key residues of WC-1 and WC-2 cycles in a circadian manner. (Left) Western Blotting (WB) was performed using protein from C-terminally V5 tagged *wc-1^1-970A&996-1167A^*grown at 25 °C and isolated at 4-hr intervals across 24 hrs following a L-D transfer. 15 μg of total protein was loaded per lane in a regular Tris-Acetate SDS-PAGE gel or a Phos-tag gel as indicated. (Right) Quantifications of WB came from three biological replicates and were plotted with x- and y-axes representing time in hrs and relative band intensities in arbitrary units. “p”, phosphorylated isoforms; “deP”, dephosphorylated isoforms. (**B**) WC-2 ChIP as a proxy for WCC DNA binding at the *frq* promoter in the dark (from (35)). Samples were cultured for indicated hrs in the dark, formaldehyde-crosslinked, and harvested; ChIP with *Neurospora* samples was done with WC-2-specific antibody. (**C**) Phosphorylation changes at S971, S990, and S433 in *wc-1^1-970A&996-1167A^* over 48 hrs. (Left) WC-1^1-970A&996-1167A^ (C-terminally V5 tagged) was immunoprecipitated by V5 antibody, and WB was performed with indicated antibodies. (Right) Bands from three independent experiments were quantified, averaged, and plotted with error bars representing +/- SEM.

To validate the Phos-tag result independently and also to track individual phosphorylation events, we raised custom antibodies against phosphorylated and unphosphorylated S971, a key phosphorylatable residue for the FFC-induced repression of WCC (28), and phosphorylated S990, a representative site for the clustered phosphorylations of S988-S995 (28, 50, 51) on WC-1, as well as phosphorylated S433 of WC-2 (17, 52), another critical phosphosite for WCC repression; all these phosphoevents are highly penetrant in the WCC population (28, 50, 52). To validate the custom antibodies against the phosphorylated- or unphosphorylated-specific residues on WC-1 and WC-2, WC-1 (V5 tagged), along with its associated WC-2, was first immunoprecipitated with V5 antibody and then treated with or without phosphatase. All the affinity-purified/depleted antibodies specifically detected their target residues with or without a phosphorylation modification (Figs. S2A and S2B). To quantify the Western Blot (WB) data from replicate studies, band intensities corresponding to unphosphorylated WC-1[S971] or phosphorylated WC-1[pS433] residues were normalized to total WC-1 or WC-2 from the same times over two circadian cycles in darkness (Fig. 2C, right-hand panels).

Consistent with the data from the Phos-tag analysis of the grouped phosphoevents (Fig. 2A), the relative level of unphosphorylated S971 increased prominently from ∼CT22 (DD8, DD32) with a peak at ∼CT2 (DD12, DD36) and declined thereafter in both cycles in darkness (Fig. 2C). Assessing small changes on a large background, levels of phosphorylation of WC-1[S971&S988-S995], bulk WC-2 (Fig. S3A), or S971 or S990 (Fig. S3B), did not display robust rhythms, although bulk WC-2 phosphorylation appears to cycle; a careful reading of data from Schafmeier *et al.* (25) reveals phosphorylation changes in bulk WC-2 consistent with this. However, as strains bearing just one phosphorylatable residue on WC-1 remain weakly rhythmic in a WT *wc-2* background (28) (Fig. S4A), it is possible to follow changes in single residues in appropriately engineered strains; thus, rhythms in the phosphorylation status of just WC-1 S971 or S990 can be followed, suggesting that dephosphorylation events are not strictly dependent on one another (Fig. S4B). Collectively, the data show that the appearance of unphosphorylated WCC correlated with its ability to bind to the *C-box* and activate *frq* expression (as predicted (17, 25, 44, 45, 53)), and also that the transcriptionally active WCC driving the oscillator represents just a small portion of total WCC, a portion that was predicted (25) but biochemically invisible unless it was specifically targeted as a population distinct from the total. This active unphosphorylated WCC must be derived from either new synthesis or dephosphorylation of the existing WCC, but first we wanted to know whether the events yielding phosphorylated inactive WCC were due mostly to FCC or reflected the overall cellular pools of CK1.

### CK1 in the FFC is involved in phosphorylation of S971 and S433

CK1 has been shown to play a crucial role in FFC-mediated phosphorylation of WCC (12, 17, 18, 54), and the strength of FRQ-CK1 steady-state interaction has been proposed to influence period length and temperature compensation (54, 55). Because CK-1a is essential for *Neurospora* (12), to assess the role of CK1 in mediating phosphorylation at the representative residues of WC-1 and WC-2, we used three documented *frq* mutants, *frq*^Δ*435-558*^, *frq^LLCN488-491AAAA^*, and *frq^QLH493-496AAA^* (12, 54), in which the FRQ and CK-1 interaction is severely impaired or even eliminated. We tested whether CK1 is involved in phosphorylation of WC-1 and WC-2 at the sites of S971 and S433 respectively.

Compatible with their performance on race tubes (12, 54), *frq*^Δ*435-558*^, *frq^LLCN488-491AAAA^*, and *frq^QLH493-496AAA^*were all arrhythmic in the *C-box-luc*-reporter assay (Fig. 3A). Interestingly, the amount of unphosphorylated WC-1[S971] was remarkably elevated while phosphorylation of WC-2[S433] dropped noticeably in the three *frq* mutants (Fig. 3B), suggesting that FFC either directly phosphorylates the two sites of WCC or achieves this indirectly through other kinases. Prior investigations have implicated CK1 in phosphorylating FRQ and governing the overall phosphorylation status of WCC (12, 17, 18, 21, 54). The data here link FRQ-bound CK1 in the FFC with the crucial phosphorylation events on WCC that lead to circadian feedback repression.

**Fig. 3.**
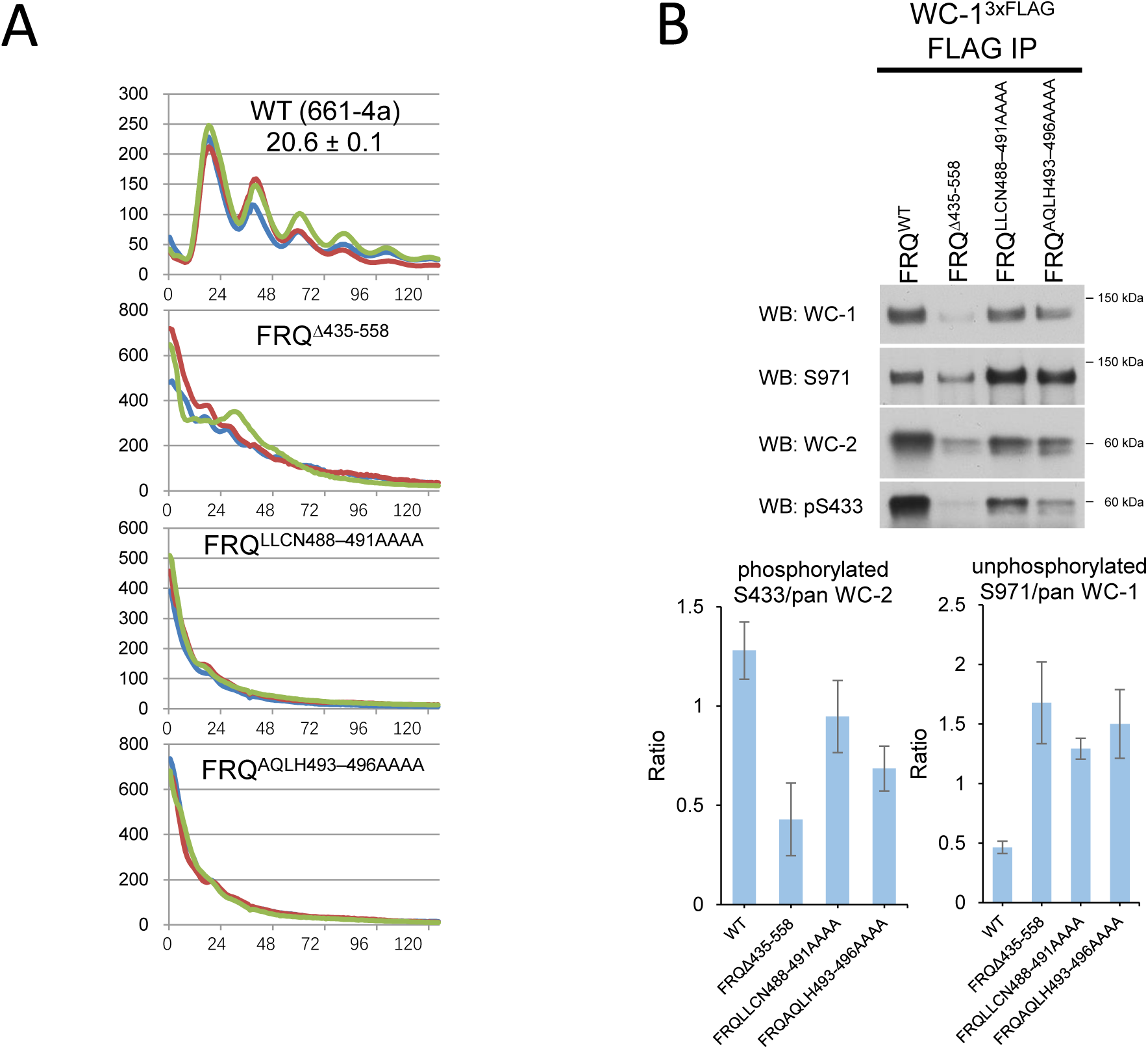
Disruption of CK1-FRQ interaction promotes the accumulation of unphosphorylated S971 of WC-1 but downregulates the abundance of phosphorylated S433 of WC-2. (**A**) Luciferase reporter assay in *frq*^Δ*435-558*^, *frq^LLCN488-491AAAA^*, and *frq^AQLH493-496AAAA^* confirming loss of rhythmicity in mutant strains compromised in FRQ-CK1 interactions. Strains grown at 25 °C in the light overnight were transferred to the dark at 25 °C for circadian clock synchronization, and bioluminescence signals were followed every hour with a CCD (charge-coupled device) camera. Three replicates (in three different colors) were plotted with the x- and y-axes representing time (in hours) and signal intensity (in arbitrary units), respectively. The *frq* mutants were targeted to the *frq* native locus with a tandem V5 and 6 x histidine (V5H6) tag at their C-termini, while WC-1 bore a 3xFLAG tag at its C-terminus. Strain 661-4a (*ras-1^bd^*, *A*, *his-3*::*C-box*-driven *luciferase*) served as WT in the assay. (**B**) (Top) Relative levels of unphosphorylated S971 and phosphorylated S433 in the *frq* mutants vs. WT. WC-1 (3xFLAG tagged) was immunoprecipitated by FLAG antibody, and WB was carried out with antibodies indicated on the left of the blots. (Bottom) Quantifications of bands from three independent experiments with error bars representing +/- SEM.

### Robust dephosphorylation of WC-1 at S971 regenerates active WCC and proceeds in the absence of de novo translation

To probe the source of unphosphorylated active WCC appearing at CT22 (DD8) (Fig. 2 and Fig. S4), the native promoter of *wc-1* was replaced by the regulatable *qa-2* promoter, a commonly used strategy to control gene expression in *Neurospora* (13); the strain also bears the *C-box*-*luc* reporter (23). In this strain synthesis of WC-1 is dependent on the inducer quinic acid (QA); the background level of expression without QA is so low that it cannot sustain rhythmicity (33) and the induced level is comparable to that seen in WT cultures (14, 33). In the first experiment, cultures were grown in the light and *wc-1* was induced by adding 10^-2^ M QA (Fig. 4A). The following morning, the *Neurospora* tissue was washed thoroughly to remove QA and thus stop *wc-1* induction, transferred to QA-minus medium, immediately moved to the dark, and sampled every 4 hrs to probe relative changes in the key phosphorylation events at the residues, S971 and S433. To prevent nonspecific binding and reduce background for better quantification in WB, and to enrich WC-1 and its bound WC-2, WC-1 (V5-tagged) was isolated from cell lysates by V5 immunoprecipitation prior to WB. First, it is plain that when the inducer QA is washed out, WC-1 is not stable but decays with a half-life of about 8 hrs indicating that new synthesis of WC-1 is required to maintain the levels seen in WT in Fig. 2A (Fig. 4A); the levels of pS971 and pS990 parallel this decrease, reflecting the total pool of WC-1. However, in contrast to this, after QA removal the relative level of unphosphorylated S971 transiently increased (Fig. 4A), consistent with what was observed from the time-course analysis of the same residue (Fig. 2C). Phosphorylation of WC-2 S433 followed the same trend as WC-1 pS971 and pS990, opposite to that of S971 in the dark (Fig. 4A). These data with those of Fig. 2 are consistent with vigorous dephosphorylation of clock-relevant residues on WCC as predicted (25). It is known, however, that there is a low level of background transcription from the *qa-2* promoter even in the absence of the inducer, QA (23), so to further confirm that the unphosphorylated WCC seen after QA washout does not come from new expression, the experiment in Fig. 4A was repeated with the addition of cycloheximide (CHX) at the time of QA withdrawal (as indicated in Fig. 4B). The patterns of WC-1 decay and dephosphorylation were similar to that seen from the CHX-free samples (Fig. 4A), although WC-1 appeared to be slightly stabilized (perhaps reflecting absence of activity-mediated turnover (56)). Again, levels of S971 rose despite decay of total WC-1, peaking similarly at DD8 and then dropping at DD12 (Fig. 4B). These data demonstrate that clock-relevant sites on WCC such as S971 can be aggressively dephosphorylated in the dark independent of new translation, consistent with prior speculation (25, 44, 45, 53), and that reactivation of WCC arising from this dephosphorylation could itself be sufficient for persistence of rhythmicity.

**Fig. 4.**
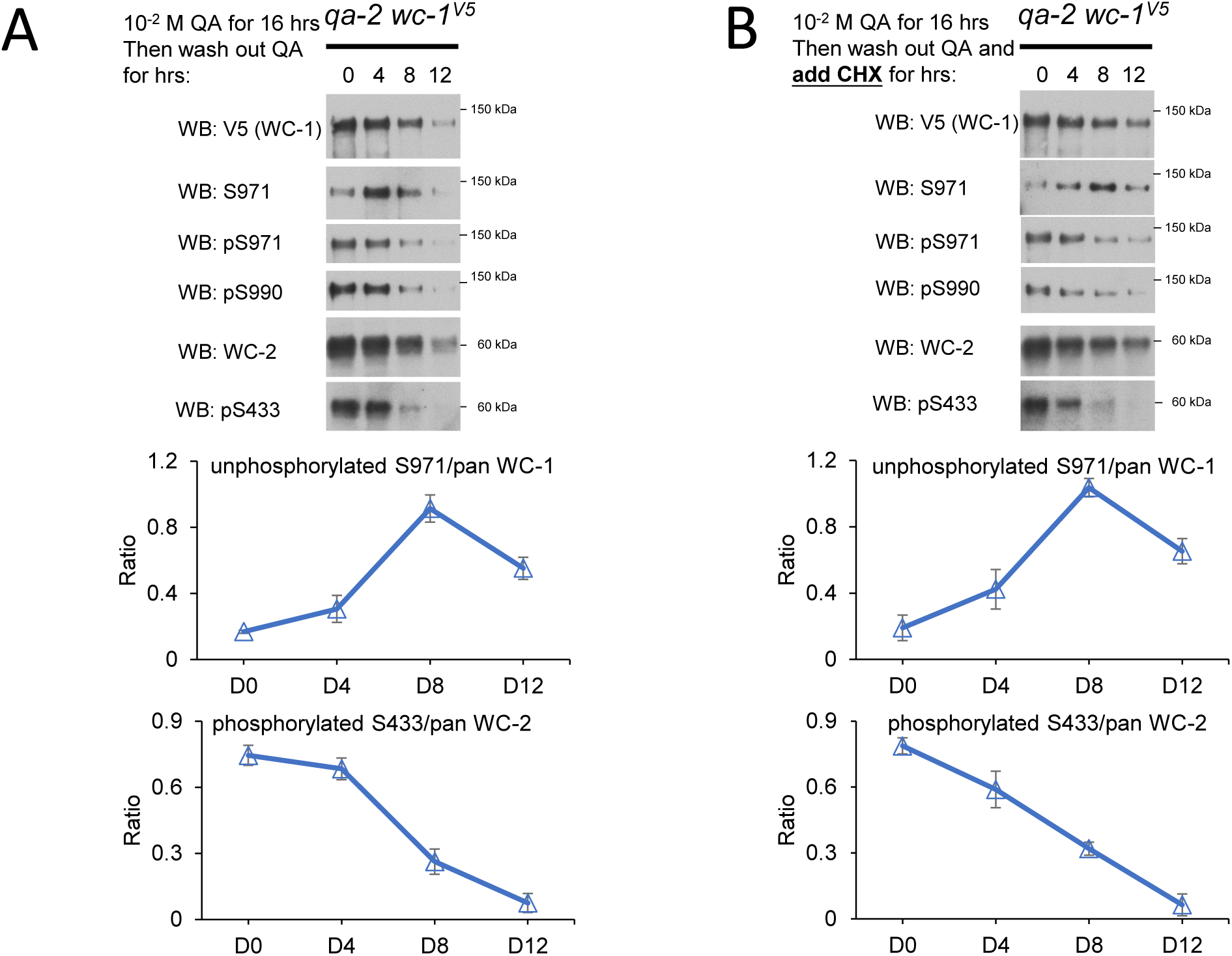
Dephosphorylation of WCC proceeds in the absence of new WC-1 expression. (**A**) (Top) *qa-2*-driven *wc-1^V5^* was grown in liquid culture medium (LCM) containing 0.1% glucose and 10^-2^ M quinic acid (QA) in the light at 25 °C overnight. Cultures were thoroughly washed, placed in 0.1% glucose LCM without QA and immediately transferred to the dark at 25 °C, and were harvested at 4, 8, or 12 hrs following the dark transfer. Protein was isolated and V5 IP (for WC-1^V5^) was performed to reduce nonspecific binding by phospho- or unphospho-specific antibodies, WB was conducted with indicated antibodies, and representative results are shown. (Bottom) Bands from three replicates were quantified, averaged, ratios calculated as shown, and plotted with error bars representing SEM. (**B**) The same experiment as in (A) was carried out with the exception of adding cycloheximide (CHX) at the final concentration of 40 µg/mL to the culture prior to the dark transfer. Quantification and plotting of WB in (A) and (B) were derived from three independent experiments.

### A cycle of dephosphorylation without new WC-1 synthesis is sufficient to operate the core oscillator

Given that termination of induction plus inhibition of translation did not prevent dephosphorylation of WC-1 at S971 (Fig. 4), we asked whether the core oscillator can persist with only dephosphorylation of existing WCC. To this end, the *qa-2:wc-1* strain was cultured in QA (10^-2^ M)-containing medium overnight, and the following morning the treated tissue was split; half was washed free of QA and cultured in medium without QA while the other was kept with QA (10^-2^ M). Both cultures were immediately moved to darkness, and rhythmicity was followed using the *frq C-box*-*luc* reporter. Another third parallel culture grown without QA throughout served as a control for background signals generated by the residual WC-1 expressed from the slightly leaky *qa-2* promoter. Cultures grown without QA showed low WCC activity (based on luciferase activity) and could be considered arrhythmic although a delayed shoulder of activity appeared in some cultures (Fig. 5A, bottom). These data suggest that the extremely low level of WC-1 from leaky expression of the promoter was not sufficient to support rhythmicity, consistent with prior observations (14, 33). Interestingly, however, the QA-induced/washed sample remained rhythmic for 3 days and the culture exposed to QA throughout for 4-5 days, both with identical ∼21 hrs period lengths (Fig. 5A). The same set of treatments as in (Fig. 5A) were repeated using a new reporter strain, *qa-2:wc-1^1-970A&996-1167A^*, in which only the six residues on WC-1 key to the core clock (28) remain phosphorylatable while all other phosphorylatable Ser and Thr in WC-1 were mutated to alanines. The rhythm now persists through three to four cycles of reactivation, the same number as the continually induced culture (Fig. 5B), and WC-1[1-970A&996-1167A] even appears to be slightly more stable than the WT (Fig. 5C), consistent with the cues driving proteasome-mediated turnover of active transcription factors (27) being among the mutated phosphosites (28). Reflecting this enhanced stability of WC-1[1-970A&996-1167A] it could be noted that even when *qa-2:wc-1^1-970A&996-1167A^* was cultured without QA throughout, it still displayed one delayed, severely dampened long period cycle that might be generated from residual WC-1, the long period being consistent with severely reduced WC-1 (33). In any case, given that rhythms can persist for at least 4 cycles in the strain *wc-1^1-970A&996-1167A^*, *wc-2^15pA^* that has only the six clock-relevant phosphorylatable residues in the WC-1 (28), these data suggest that new synthesis of WC-1 is not required for a rhythm sustained over several days nor for period-length determination of the core clock, and that derepression of the WCC via dephosphorylation alone may be sufficient for sustained rhythmicity. New WCC expression is eventually needed as existing WCC, especially WC-1, is eliminated as a part of the transcription-induced phosphorylation, ubiquitination, and degradation common to circadian and other transcriptional activators (12, 14, 25, 56, 57).

**Fig. 5.**
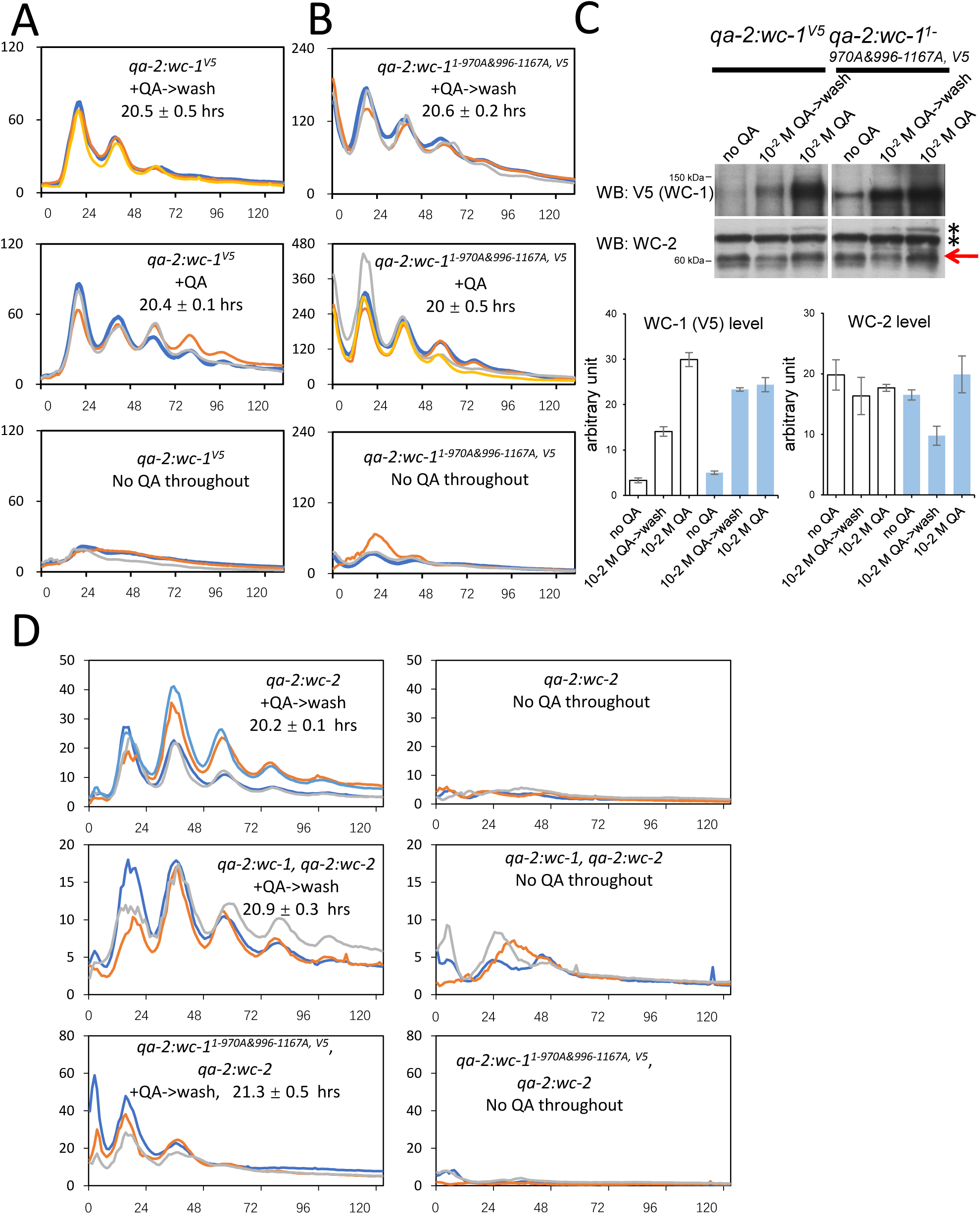
Circadian rhythms persist for several days in the absence of substantive new WCC synthesis. (**A**) Luciferase analysis of *qa-2*-driven *wc-1^V5^* under different induction conditions. The strain was cultured in LCM (0.1% glucose) with 10^-2^ M QA at 30 °C in the light. The following morning the tissue was washed thoroughly, split into two pieces, and each inoculated in LCM (0.1% glucose) in the absence (upper) or presence (middle) of 10^-2^ M QA in the light at 25 °C overnight, after which the cultures were moved to the dark for tracking bioluminescence signals in the dark at 25 °C. Bottom, another culture was treated in the same way as the other two but never encountered QA. (**B**) Luciferase analysis of *qa-2*-driven *wc-1^1-970A&996-1167A, V5^*. The same set of treatments as in (A) were performed for *qa-2*-driven *wc-1^1-970A&996-1167A, V5^*. Upper, QA was added to induce *wc-1^1-970A&996-1167A, V5^* overnight and then washed out, and afterwards luciferase signals were followed; middle, QA was supplied overnight, washed out, and then re-added; bottom, no QA was added to the medium throughout. (**C**) (Top) WB tracking WC-1 and WC-2 levels in strains as indicated at the top. Leftmost lanes corresponding to bottom panels in A and B: *Neurospora* tissue was cultured in LCM (0.1% glucose) without QA at 30 °C in the light and then transferred to 25 °C in the light and grown overnight. Middle and rightmost lanes: An overnight culture was grown with 10^-2^ M QA at 30 °C plus light, washed thoroughly to remove QA and split into two pieces. Half the tissue was cultured in the light at 25 °C overnight in the absence of QA (middle) while the other with QA at 10^-2^ M (right). Asterisks denoted nonspecific bands showing equal loadings in WB. All cultures were grown in petri dishes without shaking. (Bottom) Bands from three independent experiments were quantified, averaged, and plotted with error bars representing SEM. (**D**) Luciferase analysis of *qa-2*-driven *wc-2; qa-2*-driven *wc-1*, *qa-2*-driven *wc-2;* and *qa-2*-driven *wc-1^1-970A&996-1167A, V5^*, *qa-2*-driven *wc-2* under different induction conditions. Conidia from the indicated strains were inoculated into 2% LCM medium and incubated overnight at 30 °C. The following day, the medium was replaced with 0.1% agar-free race tube medium supplemented with 10⁻² M QA to induce expression of WC-1, WC-2, or both as indicated. Cultures were then incubated overnight at 25 °C under constant light (LL). The next morning, tissue plugs were excised, returned to the same QA-containing medium, and incubated for an additional 6 hours at 25 °C under constant light. The plugs were then washed with QA-free 0.1% race tube medium to remove QA and transferred to fresh QA-free, agar-free medium for an additional 2-hour incubation under constant light. Finally, the samples were moved to darkness at 25 °C for luciferase signal recording using a CCD camera. “No QA throughout” refers to a parallel set of treatments conducted without the addition of QA at any stage.

WB analysis was used to follow expression of WC-1 and WC-2 with samples undergoing the same set of QA treatments as for the luciferase assay (Figs. 5A and 5B). As expected, WC-1 in the two inducible strains was low when no QA was included in the medium (Fig. 5C), intermediate in the QA-supplemented/washed samples, and highest in samples that encountered QA all the time (Fig. 5C), and appeared slightly higher in the *wc-1^1-970A&996-1167A^* background, in all reflecting performance of WCC in maintaining circadian rhythms (Figs. 5A and 5B).

To examine whether dephosphorylation of WC-2 is sufficient for recycling WCC’s circadian activity, experiments parallel to those in Fig. 5A for WC-1 were performed for WC-2. It is known that WC-2 levels are depressed in *wc-2* QA-inducible strains grown without the inducer (14) in a manner like WC-1 in Fig. 5C, and in the cell, WC-2, is always in excess relative to WC-1 so its abundance does not serve as a primary determinant or a limiting component for the feedback loop (28, 58, 59). A *qa-2:wc-2* strain was grown in the presence of QA to induce WC-2 expression under light. QA was then withdrawn to terminate *wc-2* transcription, and the activity of a *C-box*-driven *luciferase* reporter was monitored in constant darkness. Following QA removal, luciferase rhythms persisted for at least four days (Fig. 5D), indicating that—similar to WC-1—preexisting WC-2 is sufficient to sustain circadian oscillations. Because WC-1 and WC-2 form a stable WCC complex, to determine whether the intact WCC complex could also function independently of new protein synthesis, we performed the same QA induction/withdrawal experiment in a *qa-2:wc-1*, *qa-2:wc-2* strain, in which expression of both WC proteins is driven by the *qa-2* promoter (14). This condition also produced multiple robust circadian cycles in constant darkness (Fig. 5D), demonstrating that preexisting WCC is competent to maintain rhythmic activation/repression activity without continuous synthesis. When this experiment was repeated using the *qa-2:wc-1^1-970A&996-1167A^*, *qa-2:wc-2* strain, two full circadian cycles were observed (Fig. 5D), suggesting that this mutant version of WC-1 may have an increased dependence on continuous WC-2 expression, but plainly establishing that daily new synthesis of WCC is not needed for rhythmicity. Taken together, our data support that existing positive element complexes in the cell are sufficient to sustain circadian rhythms in the dark through repeated cycles of dephosphorylation/activation and rephosphorylation/repression.

## Discussion

The WCC is the prototypic blue light photoreceptor for the Kingdom of Fungi as well as being an exemplar of the heterodimeric transcription factors that drive circadian oscillators in fungi and animals; in *Neurospora* it mediates both circadian regulation and acute light-regulation of a host of target genes. This biology, driven by the same protein complex, is vast and a long-standing question is why all WCC-bound promoters are not similarly regulated, i.e. why every clock-regulated WCC target is not also transcriptionally light-regulated.

Part of the evidence of this confusion, perhaps, lies in the lack of consensus regarding its recognition motif which is variously reported as centered on GATC or ATCG (9, 47, 48) with a great deal of variation in the surrounding context (e.g. (35)). As the *C-box* is weakly light-responsive (9, 47), this ambiguity may now be understood as the confounding of the two promoter types, *C-box*-like and *pLRE*-like, that have some similar and some distinct characteristics but are bound by the same transcription factor complex under different conditions. Indeed, despite the *C-box* being quiescent in constant light due to FRQ-based repression (Fig. 1), many *C-box*-like promoters have been included in lists of acutely light-responsive genes (e.g. (47, 48)); more recent biochemical analyses have identified differences in that the light-activated WCC dimer is structurally and functionally distinct from the WCC monomer in the dark: DNA binding for light-induction via *pLRE* of *frq* only needs the Zinc finger (ZnF) DNA binding domain of WC-2 while circadian DNA binding of WCC to *C-box* requires the ZnF and nearby regions of WC-2 as well as the ZnF and DBD (defective in binding DNA) motif of WC-1 (60). This is consistent with the observation that FFC-promoted phosphorylation of WC-1 does not inhibit the WCC dimer binding to *pLRE*. This distinction, now underscored by the division of genome-wide sites into the two classes, makes understanding the structural basis within the WCC for this distinction a worthy goal.

The best studied WCC-responsive promoter is that of *frq* where WCC binds at both the *C-box* and *pLRE* (9, 10). The two sets of unexpected observations reported here stem from trying to understand the importance of phosphorylation in regulating the binding activity of the WCC to these sites. The first observation is that in the light the FFC acts to remove WCC from the *C-box* but not from the *pLRE*, giving rise to the two classes of WCC-bound promoters. This finding provides a role in the light for FRQ and the FFC, a protein and complex whose roles in the cell have previously been confined to circadian regulation in the dark. Accepting that WCC may be just one of many TFs impacting expression of its targets, generalizations may still be possible as the two classes of WCC-regulated promoters have distinct behaviors: *pLRE*-like promoters display high activity at dusk and are relatively inactive at night whereas *C-box*-like have lower activity at dusk that increases through the night and decreases with the arrival of dawn. It is not difficult to imagine a potential selective advantage for two such temporally distinct and differentially light-responsive promoters.

The second unexpected observation, derived from trying to understand the first, is that during the circadian day essentially the whole pool of WC-1 is fully phosphorylated at clock-essential sites; following from this is realization of the importance of dephosphorylation, and not de novo synthesis, of WC-1 for clock function. WCC undergoes FFC-promoted phosphorylation at over 95 sites, largely the direct result of FCC-mediated CK1 activity (Fig. 4), among which only a few have been found to be necessary for circadian repression or regulating transcriptional output *ccg*–circadian amplitude. Although the causes of WCC phosphorylation/inactivation are becoming understood, prior work suggested (e.g., (12, 25)) but could not fully address the source and timing of active hypophosphorylated (or unphosphorylated) WCC, especially the central positive arm protein WC-1. Surprisingly, the pool of WC-1, even when its induction is disallowed, remains able to support several circadian cycles; plainly, de novo synthesis of WCC, especially WC-1, would be needed eventually as WC-1 is consumed when it is transcriptionally active (14, 25, 28, 51), but daily synthesis of WCC is not needed for circadian function.

Biochemical analysis has revealed that only a small percentage of FFC and WCC interact with each other in cells and do not form abundant stable complexes in the process of feedback repression (25, 61); most WCC resides in the nucleus (25) while most FFC is found in the cytoplasm (25, 61). These observations together imply transient but dynamic interactions between FFC and WCC, probably more akin to the recognition between an enzyme and its substrate (25, 61). This dynamic association enables FFC to efficiently maintain the majority of WCC in a hyperphosphorylated/inactive state, eventually leading to inactivation of the *frq* promoter; when FFC affinity for WCC is depressed by CK1-mediated FRQ phosphorylation, a small share of WCC can evade the control of FFC to be dephosphorylated, regaining its DNA binding capability (Fig. 6) and ability to rapidly drive *frq* expression. As the resulting non- or lightly phosphorylated new and functional FRQ (23), the rate-limiting factor in FFC, increases in abundance, FFC becomes activated again in promoting phosphorylation of WCC to abrogate DNA-binding and terminate *frq* expression. Interestingly, circadian regulations for FFC and WCC activities share some fundamental mechanistic basis: Although new proteins must be provided to replace those lost to turnover, coupled daily synthesis and degradation are not required for either complex to function in circadian cycles (23, 33), and multisite phosphorylation events determine and modulate their activities (17, 21, 22, 28, 62, 63); this may explain why their activating and repressing phases over circadian cycles last so long relative to proteins working in other signaling pathways, such as WCC acting in light-responses of *Neurospora* (48).

**Fig. 6.**
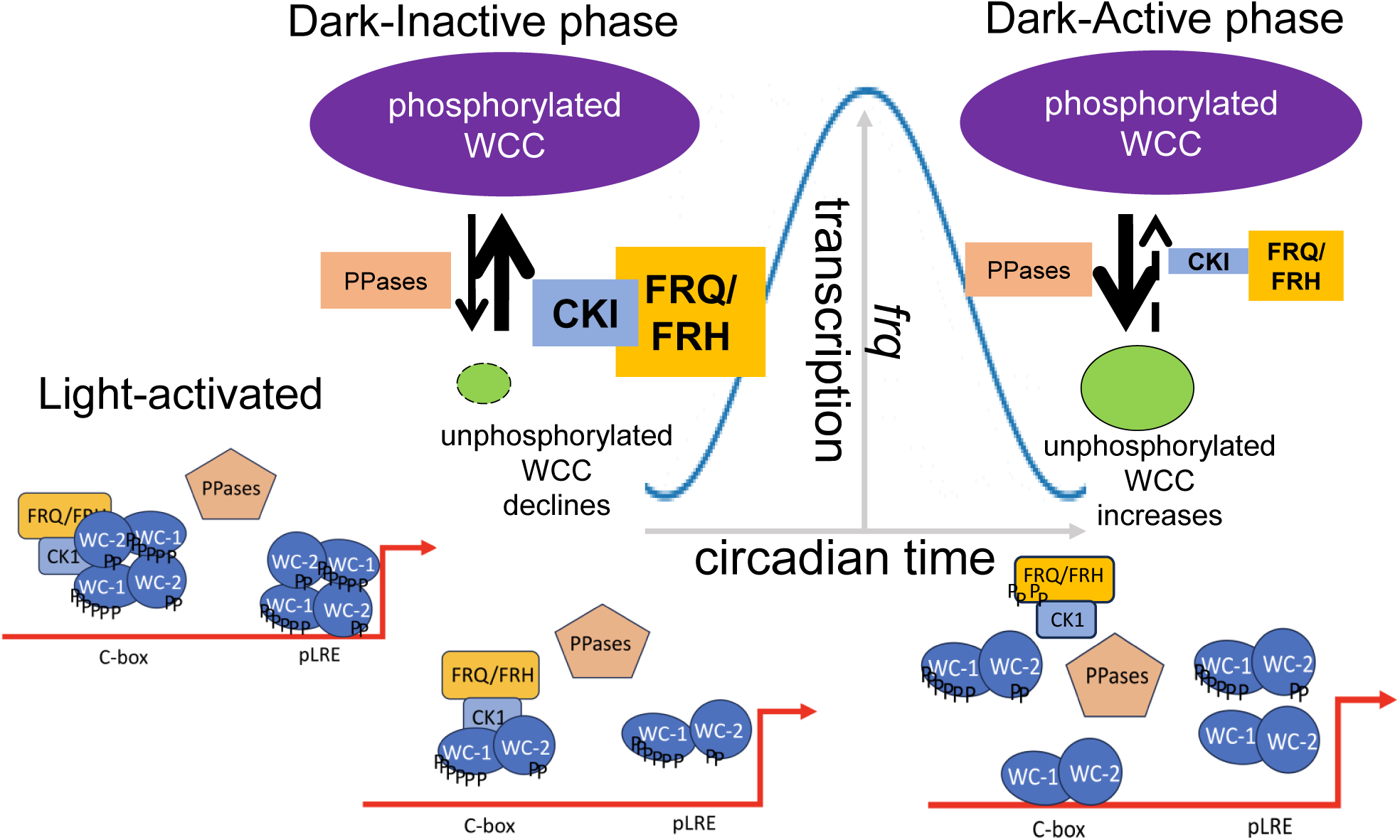
Working model showing how the overall activity of WCC in *frq* transcription is controlled by FFC and a phosphatase(s). Promoter elements and relevant proteins acting on the *frq* locus are marked. In the light, FRQ is continually expressed due to dimeric WCC action on the *pLRE*, and sites on WCC essential for binding to the *C-box* are phosphorylated maximally due to high FRQ abundance/activity, abrogating WCC binding to the *C-box*. When the light disappears (Dark-Inactive phase), new FRQ synthesis stops but existing light-induced FRQ ensures FFC-mediated WCC phosphorylation; heterodimeric WCC cannot bind the *C-box*, and the dimer of WCCs dissociates and is unable to bind the *pLRE*. FRQ becomes phosphorylated so FFC gradually loses its capacity to promote phosphorylation of WCC while the unknown phosphatases in the cell, including but not limited to PP2A (25, 28), gain the upper hand over FFC, leading to dephosphorylation/activation of a small fraction of WCC (Dark-Active phase); WCC binds the *C-box* driving *frq* transcription. New FRQ rapidly nucleates active FFC, again promoting phosphorylation of WCC in excess of the ability of the phosphatases to remove it, leading to the removal of WCC from the *frq* promoter, termination of FRQ expression, and re-establishment of the Dark-Inactive phase.

We do not know yet what phosphatases are responsible for dephosphorylating WCC to reactivate its circadian DNA binding activity, although several findings suggest that multiple phosphatases can act together. In particular, only period modifications rather than complete loss of rhythmicity have been noted in the nonessential phosphatase-deficient strains examined, and even then the effects are not internally consistent (28, 44, 45, 64). For instance, although hyperphosphorylated WCC was noted in the mutants of both RGB-1 (affecting PP2A activity), PPH4, and CSP-6 (25, 44, 46), *rgb-1^RIP^* results in a long period clock (28), whereas loss of PPH4 and PPP-1 shortens period length (44, 45) and loss of CSP-6 has negligible effects on period, although Δ*csp-6* fails to show an overt rhythm (46).

Our findings in fungi may shed some light on mammalian clocks that share a nearly identical regulatory architecture: A heterodimer of proteins interacting via PAS domains constitute a transcription factor complex that drives expression of genes whose protein products enter a complex that recruits CK1 which, in turn, phosphorylates and thereby represses the activity of the heterodimer by reducing its ability to bind to DNA. Although expression rhythms are seen in both BMAL1 and WC-1 (1, 5, 65), constitutive expression of either protein results in an essentially WT period length (31, 33, 66, 67). In mammals, both inhibition of CK1δ and activity of PPP4 leads to hypophosphorylated Bmal1 which has greater ability to bind DNA (29, 30, 43); inhibition of CK1 lengthens period in fungi and animals and inhibition of PPP4 in U2OS cells (PPH-4 in *Neurospora*) shortens period in both systems (43, 44). A unifying mechanistic model would include a dynamic balance of phosphorylation/dephosphorylation in controlling the positive arm activity, although clock proteins may act slightly differently to perform the same tasks among circadian systems. The negative element complexes (either FRQ/FRH in *Neurospora* or Pers/Crys in mammals) recruit CK1 to repress the circadian activity of their transcription factor complexes (either WCC in *Neurospora* or Bmal1/CLOCK in mammals) via phosphorylation promotion; the positive element complexes are restored to their circadian functions through timely dephosphorylation.

We have shown here that the circadian oscillator can run for several days by creating/recycling active WCC from the larger phosphorylated/inactive pool. Similarly, we have previously described a daily cycle in the ability of FFC to interact with WCC, in which newly made unphosphorylated FRQ interacts well but old heavily phosphorylated FRQ does not (17). Phosphorylated FRQ is unstable, but we now know that FRQ stability can be uncoupled from period determination (23); it is instead the phosphorylation status of FRQ that drives the cycle of FFC/WCC interaction, and thereby drives the clock. Indeed, if unphosphorylated FRQ could be generated by rapid dephosphorylation of old FRQ instead of through de novo synthesis, the clock still ought, in theory, to cycle. Stated in this way, the classic Transcription/Translation Feedback Loop (TTFL) oscillator cycle begins to take on distinctive aspects of a “Phoscillator” wherein all the clock-relevant circadian parameters, those determining periodicity, persistence, and compensation, might be explained by cycles in phosphorylation of the key players, without a requirement for daily transcription/translation at all.

## Materials and Methods

### Neurospora strains

*Neurospora* strain 661-4a (*ras-1^bd^*, *A*, *his-3*::*C-box*-driven *luciferase*) served as WT in luciferase assays; it harbors the *C-box* DNA element from the *frq* promoter fused to the codon-optimized firefly *luciferase* gene (transcriptional fusion) at the *his-3* locus. *wc-1*, *wc-2*, and *frq* mutants were engineered using a previously reported method based on homologous recombination in yeast (68). In Δ*frq* in Fig. 1C, the ORF (from the starting codon [the 1st ATG] to the stop codon TAG) of *frq* was replaced with the *bar* gene (conferring ignite resistance) while all its promoters (*C-box*, *pLRE*, and *AS*) remain intact. All introduced point mutations were verified by Sanger sequencing at the Dartmouth Genomics Core facility with *frq*-specific primers. *Neurospora* transformation was performed using an electroporator (BTX, Model # ECM630) as previously described (settings: 1,500 V, 600 Ω, and 25 μF) (22, 60).

### Growth media

Strains in vegetative growth were cultured on complete-medium slants containing 1 x Vogel’s salts, 1.6% glycerol, 0.025% casein hydrolysate, 0.5% yeast extract, 0.5% malt extract, and 1.5% agar (69). *Neurospora* sexual crosses were carried out at 25 °C with Westergaard’s agar plates bearing 1 x Westergaard’s salts, 2% sucrose, 50 ng/mL biotin, and 1.5% agar (70). Liquid culture medium (LCM) used for culturing *Neurospora* for IP and WB contains 1 x Vogel’s, 0.5% arginine, 50 ng/mL biotin, and 2% glucose (71); for quinic acid (QA) induction experiments, glucose was reduced to 0.1%. Medium used for luciferase assays contains 1 x Vogel’s salts, 0.17% arginine, 1.5% Bacto Agar, 50 ng/mL biotin, 0.1% glucose, and 12.5 μM luciferin (GoldBio, Catalog #LUCK-2G).

### Custom antibodies against the phosphorylatable residues of WC-1 and WC-2

Custom antibodies against phosphorylated S971, phosphorylated S990, or unphosphorylated S971 of WC-1 and phosphorylated S433 of WC-2 were ordered from GenScript Biotech Corporation and raised using New Zealand rabbits. Immunogenic peptides used in rabbit injections included (KKSN[p{phosphorylated}Ser]PSHSSPLHRC)-KLH (keyhole limpet haemocyanin) conjugate for phosphorylated S990, (TGNA[pSer]PTLIKGDAGC)-KLH conjugate for phosphorylated S433, (GRV[pSer]PRTSSRGGNGC)-KLH conjugate for phosphorylated S971, and (GRVSPRTSSRGGNGC)-KLH conjugate for unphosphorylated S971. Antibodies from rabbit sera were first affinity-purified with the immunogenic peptides and then immunodepleted against the immunogenic peptides with or without a phosphorylation modification at the indicated site for unphospho- or phospho-specific antibodies respectively. All the purified antibodies as the primary antibodies in WB were used by a 1: 750 dilution (20 μL antibodies to 15 mL PBST [0.3% Tween-20]), being incubated with wet-transferred protein-coated PVDF (polyvinylidene fluoride) blots in a cold room (4 °C) overnight.

### Protein extraction, immunoprecipitation (IP), and Western blot (WB)

Protein extraction, immunoprecipitation (IP), and Western blots (WB) were performed as previously described (70). Vacuum-dehydrated *Neurospora* tissue was frozen immediately in liquid nitrogen and ground to a fine powder with a ceramic mortar and pestle. 10 mL of protein-extraction buffer (50 mM HEPES [pH 7.4], 137 mM NaCl, 10% glycerol, 0.4% NP-40) was combined with one tablet of cOmplete, Mini, EDTA-free Protease Inhibitor Cocktail (Roche, Catalog # 04693159001). The inhibitor-containing buffer was mixed with the ground powder by vortexing. The mixture was treated with repeats of vortexing (10 sec) and resting on ice (10 sec) for a total of 2 min prior to incubation on ice for another 10 min. The lysate was cleared by centrifugation at 12,000 rpm at 4 °C for 10 min.

For IP using *Neurospora* lysate, 2 mg of extracts (precleared by 12,000-rpm centrifugation at 4 °C for 10 min) were incubated with 25 μL of FLAG M2 agarose (Sigma-Aldrich, Catalog # A2220) or 25 μL of V5 agarose (Sigma-Aldrich, Catalog # A7345) followed by rotating at 4 °C for 2 hrs (72). The protein-bound agarose beads were washed twice (inverting the tube ten times per wash) each with one mL of the protein-extraction buffer bearing protease inhibitors and thereafter eluted with 100 μL of 5 × SDS sample buffer by heating at 99 °C for 5 min. Ten µL out of the 100-μL IP product were loaded per lane in a gel.

For WB, 15 μg of cleared whole cell lysate was loaded per lane in a commercial 3-8% Tris-Acetate SDS gel (1.5-mm thickness, 15-well format; Thermo Fisher Scientific, Catalog # EA03785BOX) and electrophoresed using 1 x NuPAGE Tris-Acetate SDS Running Buffer (Thermo Fisher Scientific, Catalog # LA0041). The use of custom rabbit FRQ, FRH, WC-1, and WC-2 antibodies in WB has been reported previously (14, 49, 59, 73); the application of custom phospho- or unphospho-specific antibodies is detailed in the section titled “Phosphorylation-related antibodies of WC-1 and WC-2”. Phos-tag gels (6.5% SDS-PAGE containing 20 mM Phos-tag with an acrylamide:bisacrylamide ratio of 149:1) were prepared following a previous publication (28).

### Luciferase-reporter assays

Luciferase assays were done as formerly described (74).

Ninety-six–well-opaque-well plates (0.8-mL race-tube medium in each well) were inoculated with a conidial suspension, and the inoculants were grown at 25 °C in light for 16-24 hrs to saturate the wells before being transferred to the dark at 25 °C for monitoring light production. Bioluminescence signals were tracked with a CCD (charged-coupled device) camera every hour, raw data were acquired with ImageJ and a custom macro, and period lengths were manually determined.

Raw data from three replicates are shown in figures, and time (in hours) on the x-axis was plotted against the signal intensity (arbitrary units) on the y-axis.

## Acknowledgements

This study was supported by a National Institutes of Health (NIH) grant awarded to Jay C. Dunlap (R35GM118021). We acknowledge use of stocks from the Fungal Genetics Stock Center (FGSC.net), NIAID-supported informatic resources at FungiDB (Contract HHSN75N93019C00077), and Molecular Biology facilities supported by the Dartmouth CQB COBRE (NIH NIGMS grant P20GM130454), and the Dartmouth BioMT (NIH NIGMS grant P20-GM113132).

## Author contributions

B.W. and J.C.D. designed research; B.W. and X.Y.Z performed research; J.J.L provided comments; B.W. and J.C.D. wrote the paper.

---

**Fig. S1.**
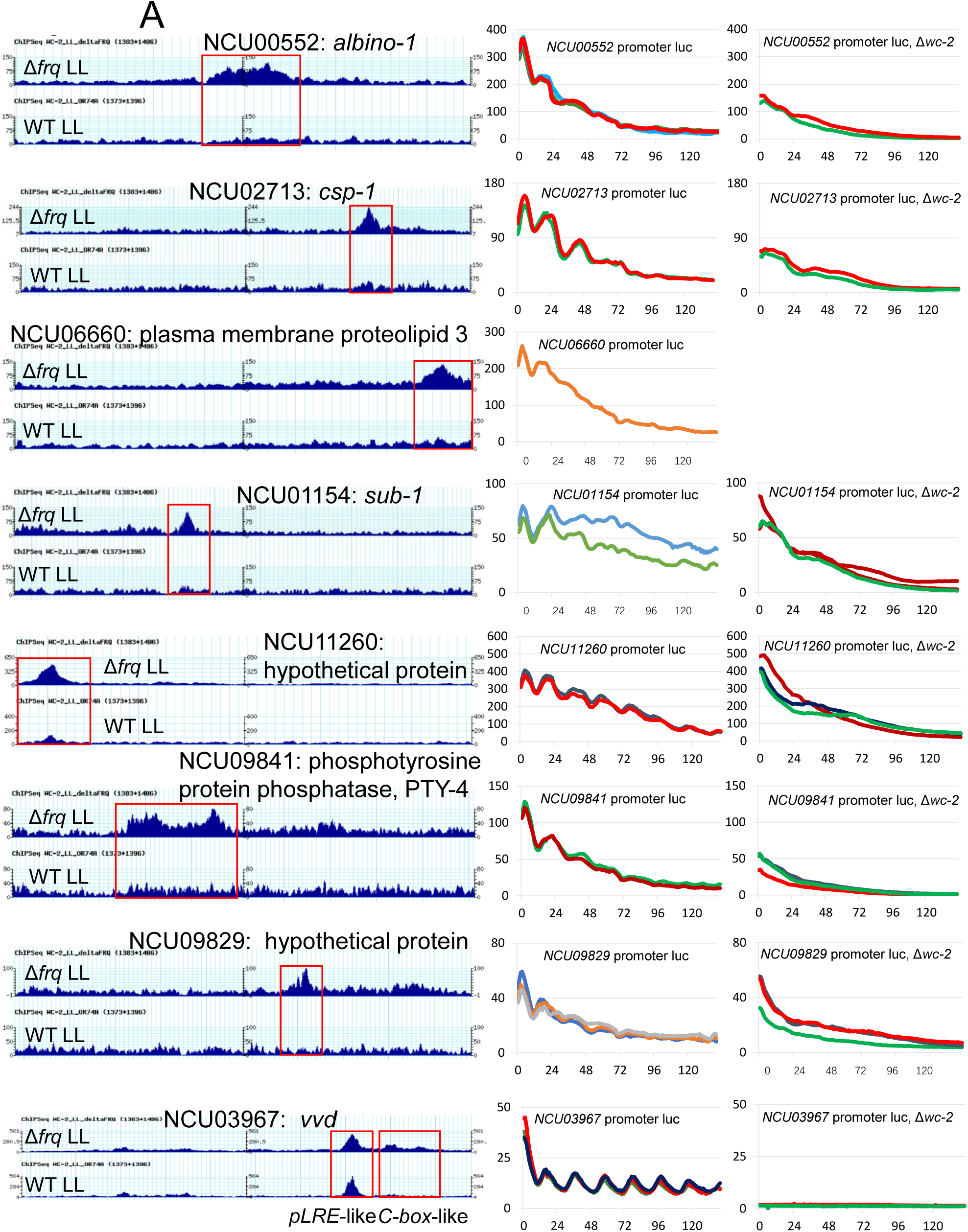
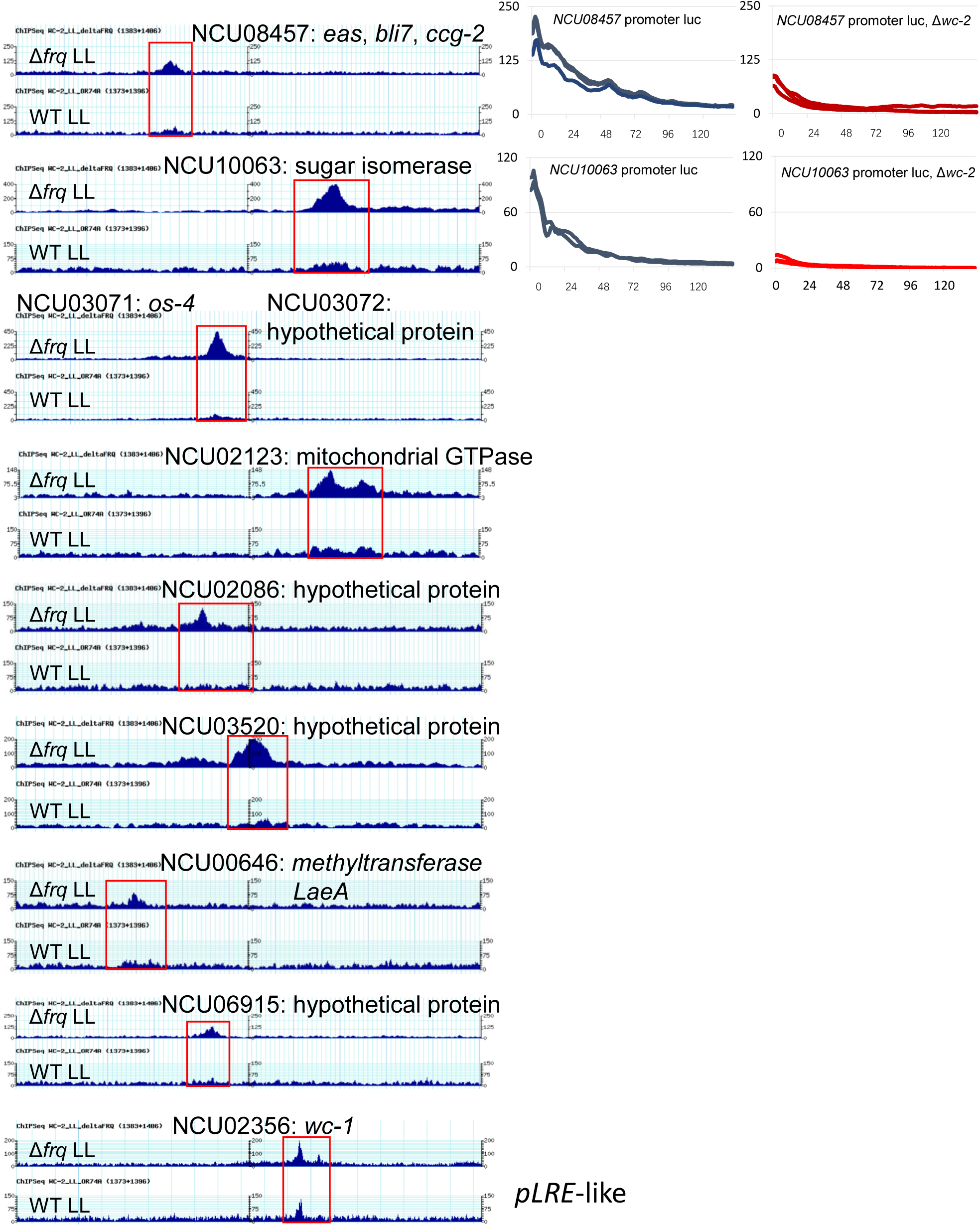
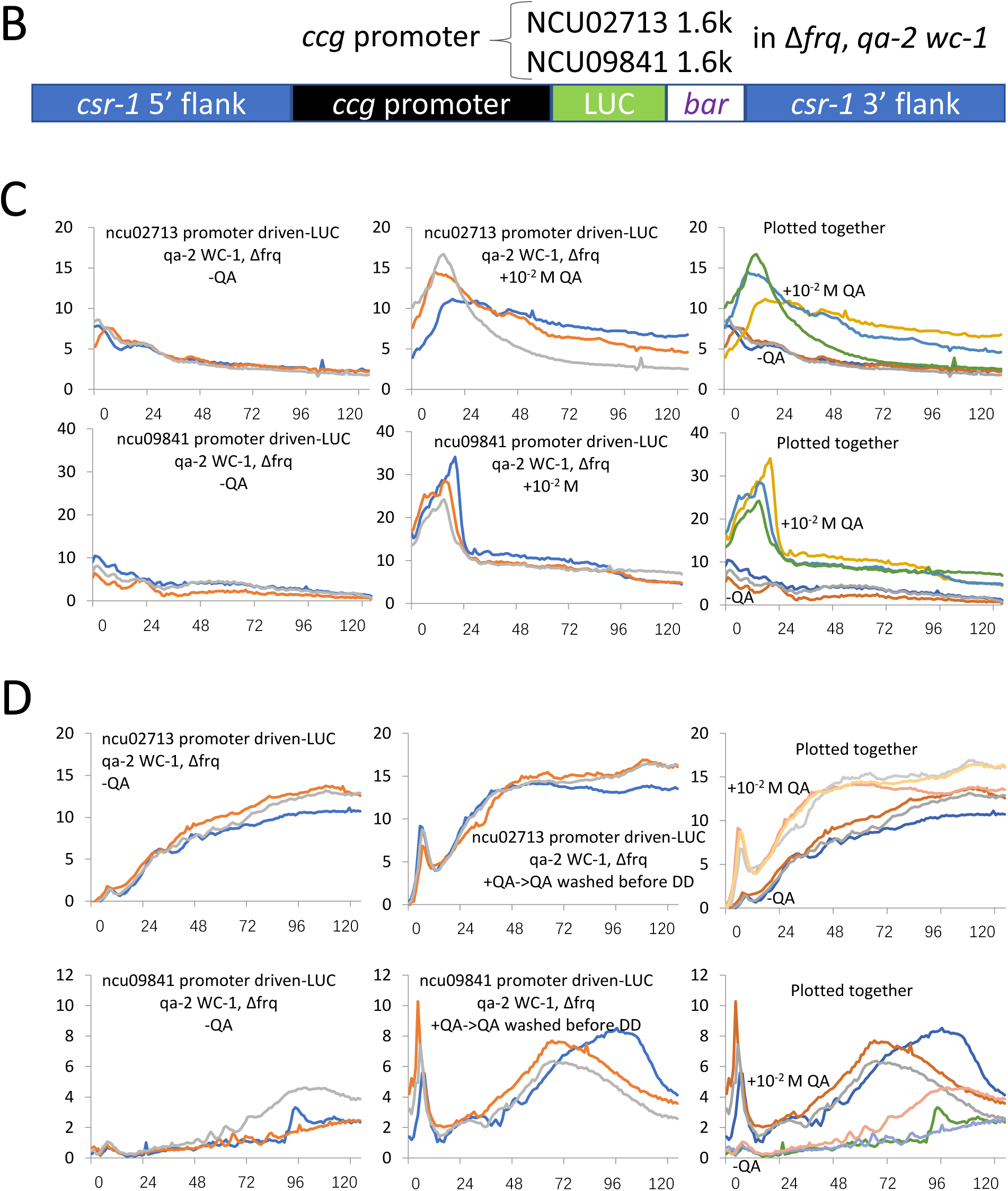
Additional examples of *C-box*-like and *pLRE*-like promoters bound by WCC. (**A**) On the left, ChIP-seq data using WC-2 antibody show WCC binding at red-boxed regions of indicated genes. For each gene, WC-2 ChIP-seq data in Δ*frq* are shown on the top while WT on the bottom; the two strains were cultured in constant light. On the right, indicated promoters were individually fused to the *luciferase* gene; the fusion constructs targeting the *csr-1* locus were transformed into either the WT (left) or Δ*wc-2* background (right) except for NCU06660 as labeled. Light production from the transgenic strains was tracked in the dark. X- and y-axes correspond to time in hrs and light intensity in arbitrary units. *C-box*-like promoters show binding of WCC in Δ*frq*, reduced binding in WT, and where examined some degree of rhythmicity in WT and a loss of rhythmicity in Δ*wc-2.* (**B**) Schematic illustration of *luciferase* constructs driven by the NCU02713 or NCU09841 promoters, integrated into the *csr-1* locus of the Δ*frq*, *qa-2:wc-1* strain. (**C**) Bioluminescence from the transgenic strains as marked was recorded in the dark, with time (in hours) on the x-axis and light intensity (in arbitrary units) on the y-axis. (**D**) Conidia from the indicated strains were inoculated into 2% LCM medium and incubated overnight at 30 °C. The following day, the medium was replaced with 0.1% race tube agar-free medium containing 10^-2^ M QA to induce WC-1 expression, and the cultures were incubated overnight at 25 °C under constant light (LL). The next morning, tissue plugs were excised, placed back into the same QA-containing medium, and incubated for an additional 6 hours at 25 °C under constant light. Subsequently, the plugs were washed with 0.1% race tube liquid medium (no QA) to remove residual QA and transferred to fresh QA-free, agar-free medium for another 2 hours of incubation under constant light. Finally, the samples were shifted to darkness at 25 °C, and luciferase signals were recorded using a camera. “−QA” refers to a parallel set of treatments carried out entirely without QA at any stage.

**Fig. S2.**
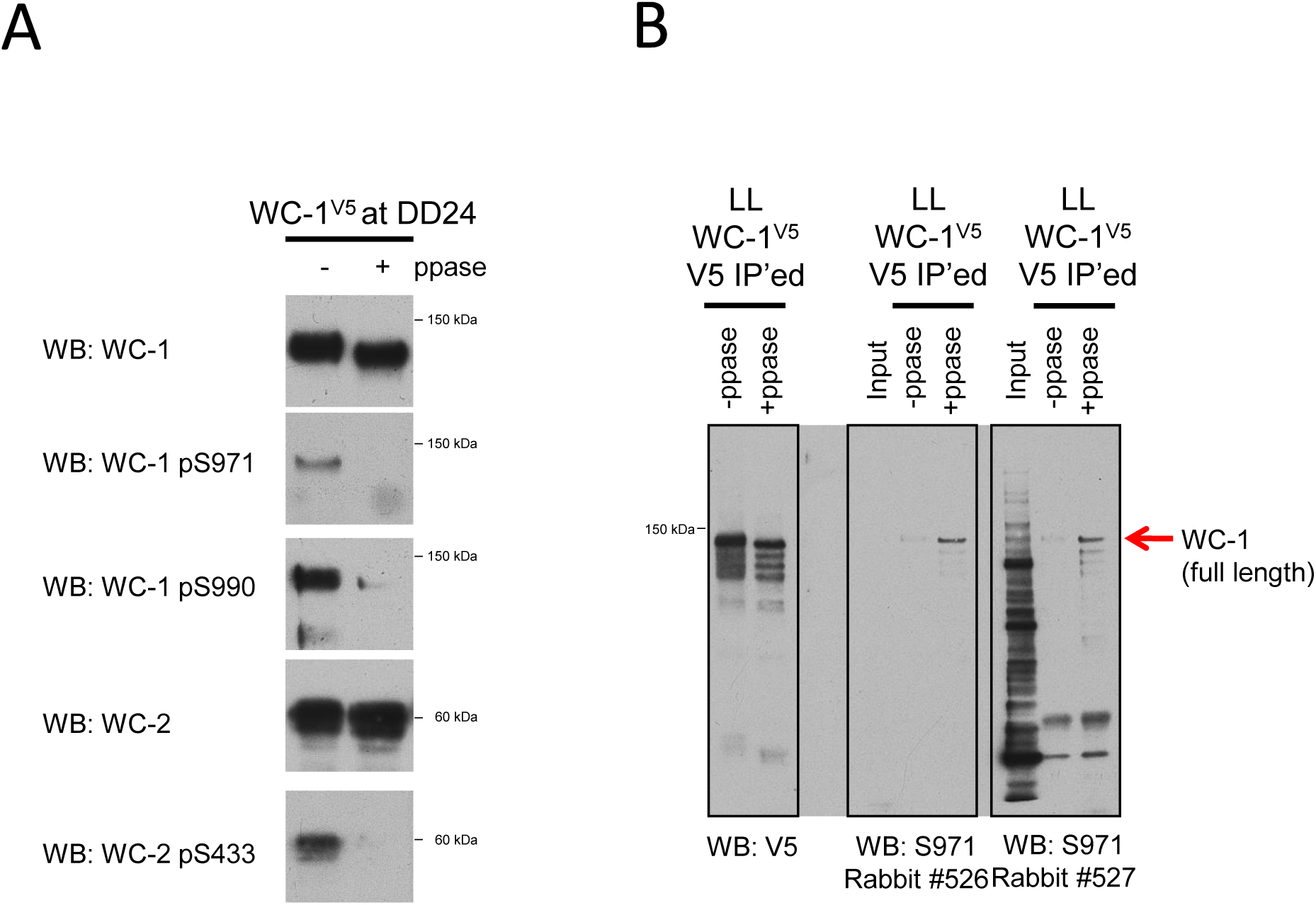
Quality control evaluation of phospho- and unphospho-specific antibodies. Custom antibodies as indicated were raised in rabbits by GenScript (see Materials and Methods for details). (**A**) *Neurospora* protein extracts were made from cultures grown for 24 hrs in darkness. WC-1^V5^ was pulled down with V5 antibody and then treated with or without Lambda phosphatase. WB was developed with indicated antibodies against phosphorylated S971 (pS971), S990 (pS990), or S433 (pS433). (**B**) Validation of unphosphorylated-specific antibody against S971 of WC-1. Cultures were grown as in (A) except under constant light (LL). Each phosphorylation-related antibody was used as a primary antibody in WB at a dilution of 1:750.

**Fig. S3.**
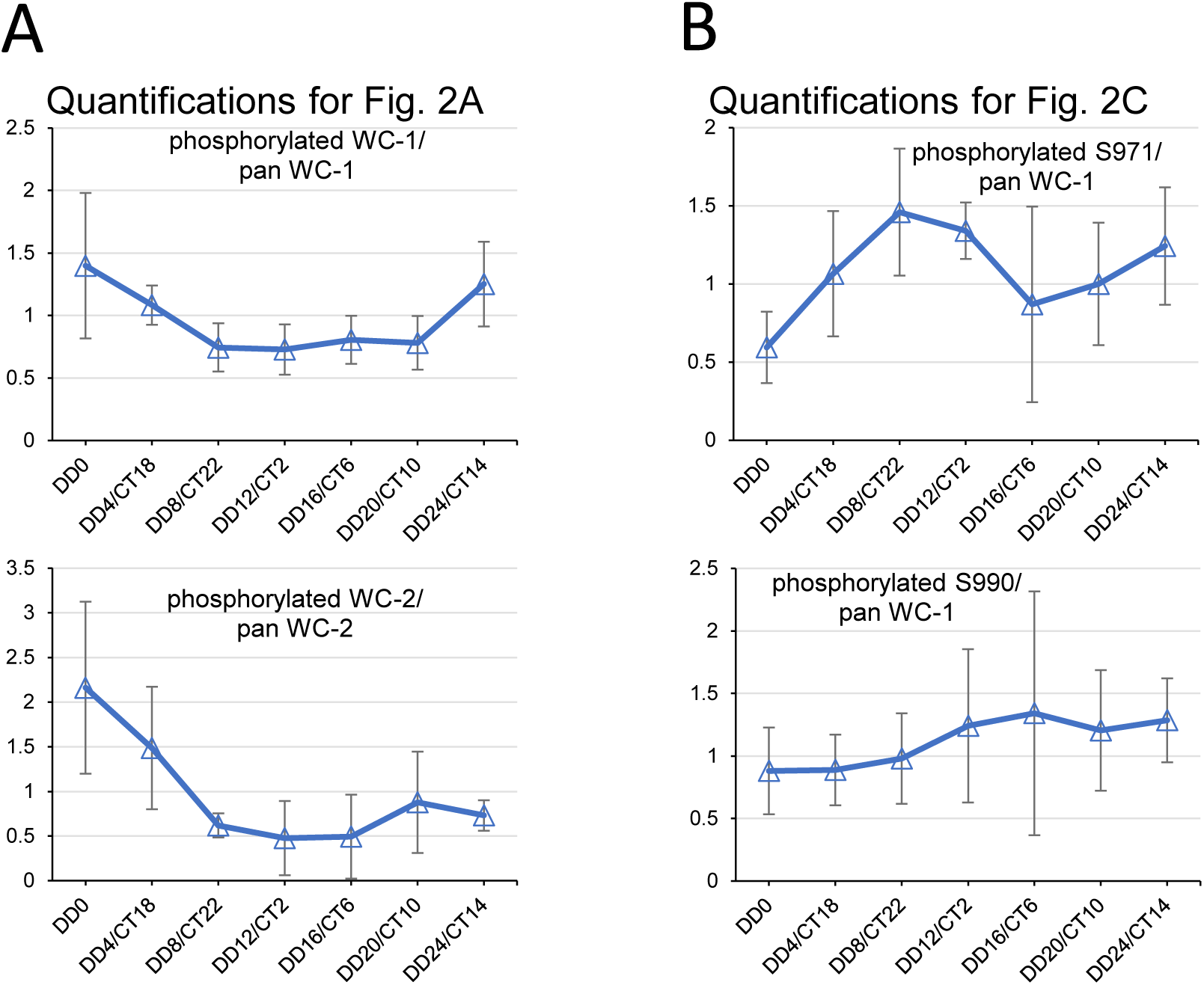
Phosphorylation of total pools of WC-1 and WC-2 is not strongly rhythmic. (**A**) Quantifications of phosphorylated WC-1 and WC-2 in Fig. 2A (upper bands in the Phos-tag gel blots with V5 and WC-2 antibodies respectively) normalized to total WC-1 and WC-2 (bands in the regular gel blots of V5 and WC-2). (**B**) Quantifications of phosphorylated S971 and S990 bands in Fig. 2C developed with phosphorylation-specific antibodies. Quantifications in (A) and (B) were both performed with three independent experiments; error bars are SEM. Statistical analyses were performed for data from two adjacent circadian times (CTs). Differences between any two adjacent CTs are considered not statistically significant (https://www.graphpad.com/quickcalcs/ttest1/?format=SEM; the two-tailed P values range from 0.2 to 1).

**Fig. S4.**
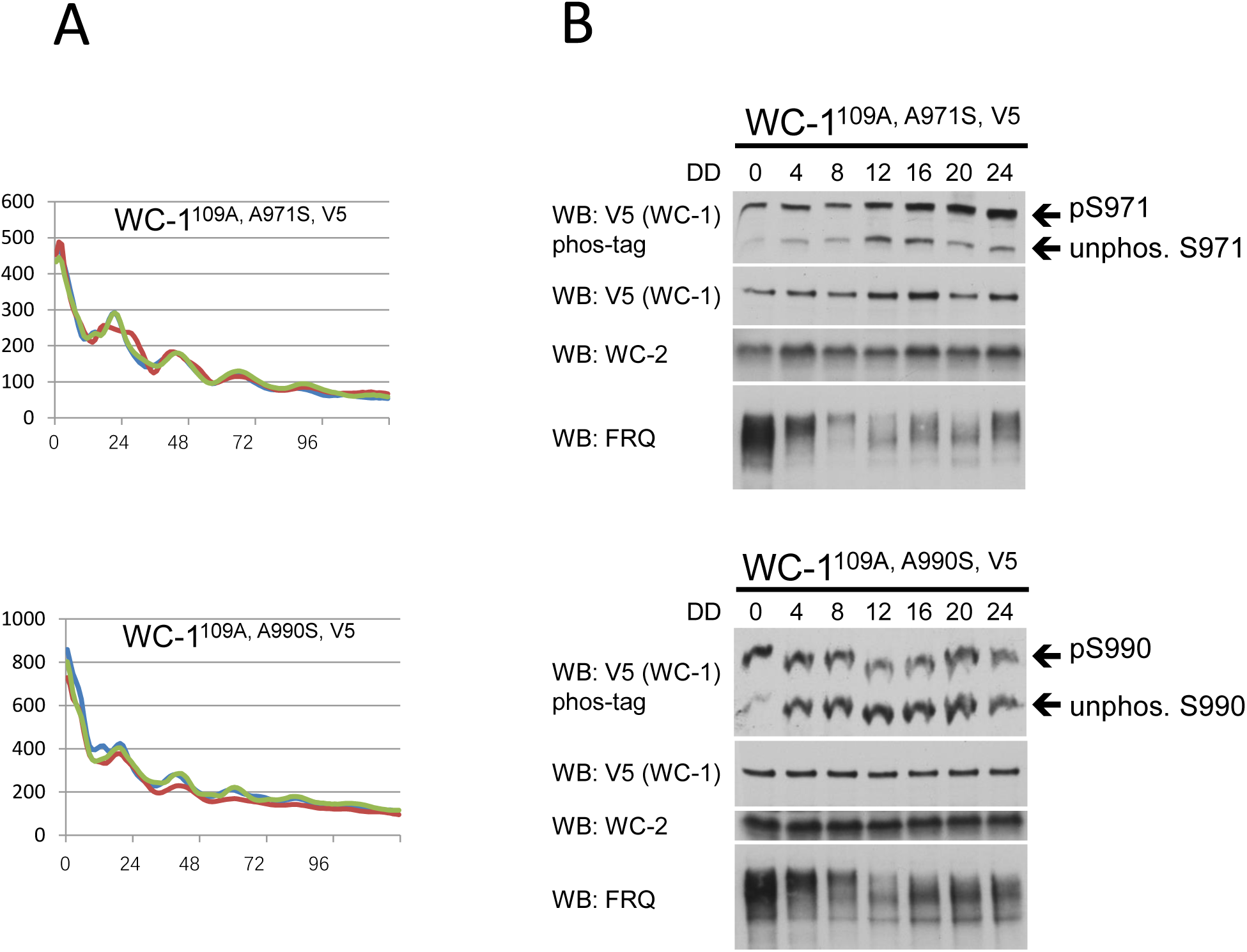
Phosphorylation status of S971 and S990 of WC-1 changes over a circadian cycle. (**A**) Luciferase assays of *wc-1^109A, A971S, V5^* (upper) or *wc-1^109A, A990S, V5^* (bottom). Clock synchronization and bioluminescence signal acquirement were both completed at 25 °C in the presence and absence of light respectively. The *C-box*-driven *luciferase* in both strains is at the *his-3* locus. WC-2 is WT in this strain, so it remains rhythmic. (**B**) Phos-tag analyses of WC-1 S971 and S990 over one day were performed for *wc-1^109A, A971S, V5^* (upper) or *wc-1^109A, A990S, V5^* (bottom). In these strains 77 known WC-1 phosphosites and 32 serines/threonines were all mutated to alanines except for S971 or S990 as indicated (28).

**Table S1.** Summary of identified *C-box*-like and *pLRE*-like promoters as evidenced by WC-2 ChIP-seq from cultures grown in the light. For reference, the classification of genes as early-light-responding or late-light-responding from Chen et al., 2009 is included. The raw WC-2 ChIP-seq data files have been deposited into the NCBI BioProject database with ID# PRJNA1122768.

| C-box like binding pattern | Gene number | Gene name | Type | Chen et al., 2009 |
| --- | --- | --- | --- | --- |
|  | NCU00552T1 | *albino-1* | C-box like | Early |
|  | NCU00646T1 | methyltransferase LaeA | C-box like |  |
|  | NCU00938T1 | hypothetical protein | C-box like |  |
|  | NCU01154T2 | *sub-1* | C-box like | Early |
|  | NCU02123T1 | mitochondrial GTPase | C-box like | NCU02124 late |
|  | NCU02264T1 | hypothetical protein | C-box like | Early |
|  | NCU02265T3 | *frq* | C-box like | Early |
|  | NCU02713T1 | *csp-1* | C-box like | Early |
|  | NCU02724T1 | HLH transcript factors | C-box like |  |
|  | NCU02755T1 | PAP2 domain | C-box like |  |
|  | NCU03071T1 | *os-4* | C-box like | Early |
|  | NCU03072T1 | hypothetical protein | C-box like | Early |
|  | NCU03520T1 | hypothetical protein | C-box like |  |
|  | NCU06593T1 | MAP kinase activator | C-box like |  |
|  | NCU06660T1 | plasma membrane proteolipid 3 | C-box like |  |
|  | NCU07024T1 | *os-2/Mak-3* | C-box like | Early |
|  | NCU07540T1 | hypothetical protein | C-box like | Early |
|  | NCU08457T1 | *eas, bli7, ccg-2* | C-box like |  |
|  | NCU08607T1 | ER Golgi intermediate compartment protein 3 | C-box like |  |
|  | NCU11260T0 | hypothetical protein | C-box like |  |
|  | NCU04005T1 | *ck-1b* | C-box like |  |
|  | NCU11290T0 | hypothetical protein | C-box like |  |
|  | NCU09841T1 | phosphotyrosine protein phosphatase | C-box like |  |
|  | NCU09829T1<br>NCU09830 | hypothetical protein | C-box like | Early |
|  | NCU02086T1 | hypothetical protein | C-box like |  |
|  | NCU08443 | Hypothetical protein (TF candidate) | C-box like |  |

| pLRE like binding pattern | Gene number | Gene name | Type | Chen et al., 2009 |
| --- | --- | --- | --- | --- |
|  | NCU09535T1 | hypothetical protein | more similar to *pLRE* | Early |
|  | NCU02356T1 | *wc-1* | *pLRE* and C-box like | Early/Late |
|  | NCU10063T1 | sugar isomerase | *pLRE* like | Early |
|  | NCU02801T1 | translocator protein | *pLRE* like | Early |
|  | NCU06915T1 | hypothetical protein | *pLRE* like | Early |
|  | NCU04832T1 | dehydropantoate reductase-1 | *pLRE* like |  |
|  | NCU00582T1 | cytochrome | *pLRE* like | Early/Late |
|  | NCU01926T1 | hypothetical protein | *pLRE* like |  |
|  | NCU02819T3 | high mobility group protein | *pLRE* like | Early/Late |
|  | NCU03967T1 | *vvd* | *pLRE* and C-box like | Early/Late |
|  | NCU07541T1 | hypothetical protein | *pLRE* like | Early |
|  | NCU07659T1 | acetate-4 | *pLRE* like |  |
|  | NCU08402T1 | zinc-binding alcohol dehydrogenase | *pLRE* like (not strong) |  |
|  | NCU08850T1 | mutagen sensitive-18 | *pLRE* like (not strong) | Early |
|  | NCU05340T1 | alkanesulfonate monooxygenase | *pLRE* like/C-box |  |
|  | NCU06483T1 | F-box and WD repeat-containing protein | *pLRE* like? |  |
|  | NCU04417T1 | hypothetical protein | Either, just peak binding | Late |
|  | NCU08161 | Histone chaperone | *pLRE* like | Early |
|  | NCU02190 | Oxysterol binding protein | *pLRE* like | Early |
|  | NCU00939 | Development and carotenogenesis control -1 | *pLRE*-like | Early |

## References

1. D. Bell-Pedersen, et al., Circadian rhythms from multiple oscillators: lessons from diverse organisms. Nat Rev Genet 6, 544–556 (2005).

2. T. Roenneberg, M. Merrow, The Circadian Clock and Human Health. Current Biology 26, R432–R443 (2016).

3. C. R. Cederroth, et al., Medicine in the Fourth Dimension. Cell Metab 30, 238–250 (2019).

4. J. Cha, M. Zhou, Y. Liu, Mechanism of the Neurospora Circadian Clock, a FREQUENCY-centric View. Biochemistry 54, 150–156 (2015).

5. C. L. Partch, Orchestration of Circadian Timing by Macromolecular Protein Assemblies. Journal of Molecular Biology 432, 3426–3448 (2020).

6. J. C. Dunlap, J. J. Loros, Making Time: Conservation of Biological Clocks from Fungi to Animals. Microbiol Spectr 5, 5.3.05 (2017).

7. C. Partch, M. Brunner, How circadian clocks keep time: the discovery of slowness. FEBS Letters 596, 1613–1614 (2022).

8. Q. He, et al., White Collar-1, a DNA Binding Transcription Factor and a Light Sensor. Science 297, 840–843 (2002).

9. A. C. Froehlich, Y. Liu, J. J. Loros, J. C. Dunlap, White Collar-1, a Circadian Blue Light Photoreceptor, Binding to the frequency Promoter. Science 297, 815–819 (2002).

10. A. C. Froehlich, J. J. Loros, J. C. Dunlap, Rhythmic binding of a WHITE COLLAR-containing complex to the frequency promoter is inhibited by FREQUENCY. Proc. Natl. Acad. Sci. U.S.A. 100, 5914–5919 (2003).

11. W. J. Belden, J. J. Loros, J. C. Dunlap, Execution of the Circadian Negative Feedback Loop in Neurospora Requires the ATP-Dependent Chromatin-Remodeling Enzyme CLOCKSWITCH. Molecular Cell 25, 587–600 (2007).

12. Q. He, et al., CKI and CKII mediate the FREQUENCY-dependent phosphorylation of the WHITE COLLAR complex to close the Neurospora circadian negative feedback loop. Genes Dev. 20, 2552–2565 (2006).

13. B. D. Aronson, K. A. Johnson, J. J. Loros, J. C. Dunlap, Negative Feedback Defining a Circadian Clock: Autoregulation of the Clock Gene frequency. Science 263, 1578–1584 (1994).

14. M. Shi, M. Collett, J. J. Loros, J. C. Dunlap, FRQ-Interacting RNA Helicase Mediates Negative and Positive Feedback in the Neurospora Circadian Clock. Genetics 184, 351–361 (2010).

15. P. Cheng, Q. He, Q. He, L. Wang, Y. Liu, Regulation of the Neurospora circadian clock by an RNA helicase. Genes Dev. 19, 234–241 (2005).

16. J. Guo, P. Cheng, Y. Liu, Functional Significance of FRH in Regulating the Phosphorylation and Stability of Neurospora Circadian Clock Protein FRQ. Journal of Biological Chemistry 285, 11508–11515 (2010).

17. C. L. Baker, A. N. Kettenbach, J. J. Loros, S. A. Gerber, J. C. Dunlap, Quantitative Proteomics Reveals a Dynamic Interactome and Phase-Specific Phosphorylation in the Neurospora Circadian Clock. Molecular Cell 34, 354–363 (2009).

18. D. Marzoll, et al., Casein kinase 1 and disordered clock proteins form functionally equivalent, phospho-based circadian modules in fungi and mammals. Proc. Natl. Acad. Sci. U.S.A. 119, e2118286119 (2022).

19. K. Vanselow, A. Kramer, Role of Phosphorylation in the Mammalian Circadian Clock. Cold Spring Harbor Symposia on Quantitative Biology 72, 167–176 (2007).

20. K. L. Ode, et al., Knockout-Rescue Embryonic Stem Cell-Derived Mouse Reveals Circadian-Period Control by Quality and Quantity of CRY1. Molecular Cell 65, 176–190 (2017).

21. C.-T. Tang, et al., Setting the pace of the Neurospora circadian clock by multiple independent FRQ phosphorylation events. Proc. Natl. Acad. Sci. U.S.A. 106, 10722–10727 (2009).

22. B. Wang, E.-L. Stevenson, J. C. Dunlap, Functional analysis of 110 phosphorylation sites on the circadian clock protein FRQ identifies clusters determining period length and temperature compensation. G3 Genes|Genomes|Genetics 13, jkac334 (2023).

23. L. F. Larrondo, C. Olivares-Yañez, C. L. Baker, J. J. Loros, J. C. Dunlap, Decoupling circadian clock protein turnover from circadian period determination. Science 347, 1257277 (2015).

24. A. Kramer, When the circadian clock becomes blind. Science 347, 476–477 (2015).

25. T. Schafmeier, et al., Transcriptional Feedback of Neurospora Circadian Clock Gene by Phosphorylation-Dependent Inactivation of Its Transcription Factor. Cell 122, 235–246 (2005).

26. C. Lee, J.-P. Etchegaray, F. R. A. Cagampang, A. S. I. Loudon, S. M. Reppert, Posttranslational Mechanisms Regulate the Mammalian Circadian Clock. Cell 107, 855–867 (2001).

27. H. Yoshitane, et al., Roles of CLOCK Phosphorylation in Suppression of E-Box-Dependent Transcription. Molecular and Cellular Biology 29, 3675–3686 (2009).

28. B. Wang, A. N. Kettenbach, X. Zhou, J. J. Loros, J. C. Dunlap, The Phospho-Code Determining Circadian Feedback Loop Closure and Output in Neurospora. Molecular Cell 74, 771–784.e3 (2019).

29. X. Cao, Y. Yang, C. P. Selby, Z. Liu, A. Sancar, Molecular mechanism of the repressive phase of the mammalian circadian clock. Proc. Natl. Acad. Sci. U.S.A. 118, e2021174118 (2021).

30. Y. An, et al., Decoupling PER phosphorylation, stability and rhythmic expression from circadian clock function by abolishing PER-CK1 interaction. Nat Commun 13, 3991 (2022).

31. A. C. Liu, et al., Redundant function of REV-ERBalpha and beta and non-essential role for Bmal1 cycling in transcriptional regulation of intracellular circadian rhythms. PLoS Genet 4, e1000023 (2008).

32. Y. Otobe, et al., Phosphorylation of DNA-binding domains of CLOCK–BMAL1 complex for PER-dependent inhibition in circadian clock of mammalian cells. Proc. Natl. Acad. Sci. U.S.A. 121, e2316858121 (2024).

33. P. Cheng, Y. Yang, Y. Liu, Interlocked feedback loops contribute to the robustness of the Neurospora circadian clock. Proc. Natl. Acad. Sci. U.S.A. 98, 7408–7413 (2001).

34. J. M. Hurley, et al., Circadian Proteomic Analysis Uncovers Mechanisms of Post-Transcriptional Regulation in Metabolic Pathways. Cell Systems 7, 613–626.e5 (2018).

35. J. M. Hurley, et al., Analysis of clock-regulated genes in Neurospora reveals widespread posttranscriptional control of metabolic potential. Proc. Natl. Acad. Sci. U.S.A. 111, 16995–17002 (2014).

36. W. Dong, et al., Systems Biology of the Clock in Neurospora crassa. PLoS ONE 3, e3105 (2008).

37. H. Cho, et al., Regulation of circadian behaviour and metabolism by REV-ERB-α and REV-ERB-β. Nature 485, 123–127 (2012).

38. Y. Fang, S. Sathyanarayanan, A. Sehgal, Post-translational regulation of the Drosophila circadian clock requires protein phosphatase 1 (PP1). Genes Dev. 21, 1506–1518 (2007).

39. S. Sathyanarayanan, X. Zheng, R. Xiao, A. Sehgal, Posttranslational Regulation of Drosophila PERIOD Protein by Protein Phosphatase 2A. Cell 116, 603–615 (2004).

40. H. Lee, et al., The period of the circadian oscillator is primarily determined by the balance between casein kinase 1 and protein phosphatase 1. Proc. Natl. Acad. Sci. U.S.A. 108, 16451–16456 (2011).

41. I. Schmutz, et al., Protein Phosphatase 1 (PP1) Is a Post-Translational Regulator of the Mammalian Circadian Clock. PLoS ONE 6, e21325 (2011).

42. C. L. Partch, K. F. Shields, C. L. Thompson, C. P. Selby, A. Sancar, Posttranslational regulation of the mammalian circadian clock by cryptochrome and protein phosphatase 5. Proc. Natl. Acad. Sci. U.S.A. 103, 10467–10472 (2006).

43. S. Klemz, et al., Protein phosphatase 4 controls circadian clock dynamics by modulating CLOCK/BMAL1 activity. Genes Dev. 35, 1161–1174 (2021).

44. Y. Yang, et al., Distinct roles for PP1 and PP2A in the Neurospora circadian clock. Genes Dev. 18, 255–260 (2004).

45. J. Cha, S.-S. Chang, G. Huang, P. Cheng, Y. Liu, Control of WHITE COLLAR localization by phosphorylation is a critical step in the circadian negative feedback process. EMBO J 27, 3246–3255 (2008).

46. X. Zhou, et al., A HAD family phosphatase CSP-6 regulates the circadian output pathway in Neurospora crassa. PLoS Genet 14, e1007192 (2018).

47. K. M. Smith, et al., Transcription Factors in Light and Circadian Clock Signaling Networks Revealed by Genomewide Mapping of Direct Targets for Neurospora White Collar Complex. Eukaryot Cell 9, 1549–1556 (2010).

48. C.-H. Chen, C. S. Ringelberg, R. H. Gross, J. C. Dunlap, J. J. Loros, Genome-wide analysis of light-inducible responses reveals hierarchical light signalling in Neurospora. EMBO J 28, 1029–1042 (2009).

49. N. Y. Garceau, Y. Liu, J. J. Loros, J. C. Dunlap, Alternative Initiation of Translation and Time-Specific Phosphorylation Yield Multiple Forms of the Essential Clock Protein FREQUENCY. Cell 89, 469–476 (1997).

50. G. Huang, et al., Protein kinase A and casein kinases mediate sequential phosphorylation events in the circadian negative feedback loop. Genes Dev. 21, 3283–3295 (2007).

51. Q. He, et al., Light-independent Phosphorylation of WHITE COLLAR-1 Regulates Its Function in the Neurospora Circadian Negative Feedback Loop. Journal of Biological Chemistry 280, 17526–17532 (2005).

52. G. Sancar, C. Sancar, M. Brunner, T. Schafmeier, Activity of the circadian transcription factor White Collar Complex is modulated by phosphorylation of SP-motifs. FEBS Letters 583, 1833–1840 (2009).

53. Q. He, Y. Liu, Molecular mechanism of light responses in Neurosporaϑ: from light-induced transcription to photoadaptation. Genes Dev. 19, 2888–2899 (2005).

54. X. Liu, et al., FRQ-CK1 interaction determines the period of circadian rhythms in Neurospora. Nat Commun 10, 4352 (2019).

55. Y. Hu, et al., FRQ-CK1 Interaction Underlies Temperature Compensation of the Neurospora Circadian Clock. mBio 12, e01425–21 (2021).

56. W. P. Tansey, Transcriptional activation: risky business. Genes Dev. 15, 1045–1050 (2001).

57. M. Stratmann, D. M. Suter, N. Molina, F. Naef, U. Schibler, Circadian Dbp Transcription Relies on Highly Dynamic BMAL1-CLOCK Interaction with E Boxes and Requires the Proteasome. Molecular Cell 48, 277–287 (2012).

58. P. Cheng, Y. Yang, K. H. Gardner, Y. Liu, PAS Domain-Mediated WC-1/WC-2 Interaction Is Essential for Maintaining the Steady-State Level of WC-1 and the Function of Both Proteins in Circadian Clock and Light Responses of Neurospora. Mol Cell Biol 22, 517–524 (2002).

59. D. L. Denault, WC-2 mediates WC-1-FRQ interaction within the PAS protein-linked circadian feedback loop of Neurospora. The EMBO Journal 20, 109–117 (2001).

60. B. Wang, X. Zhou, J. J. Loros, J. C. Dunlap, Alternative Use of DNA Binding Domains by the Neurospora White Collar Complex Dictates Circadian Regulation and Light Responses. Molecular and Cellular Biology 36, 781–793 (2016).

61. C. I. Hong, P. Ruoff, J. J. Loros, J. C. Dunlap, Closing the circadian negative feedback loop: FRQ-dependent clearance of WC-1 from the nucleus. Genes Dev. 22, 3196–3204 (2008).

62. X. Chen, et al., Differential regulation of phosphorylation, structure and stability of circadian clock protein FRQ isoforms. Journal of Biological Chemistry 104597 (2023). 10.1016/j.jbc.2023.104597.

63. Y. Liu, J. Loros, J. C. Dunlap, Phosphorylation of the Neurospora clock protein FREQUENCY determines its degradation rate and strongly influences the period length of the circadian clock. Proc. Natl. Acad. Sci. U.S.A. 97, 234–239 (2000).

64. Z. Ding, T. M. Lamb, A. Boukhris, R. Porter, D. Bell-Pedersen, Circadian Clock Control of Translation Initiation Factor eIF2α Activity Requires eIF2γ-Dependent Recruitment of Rhythmic PPP-1 Phosphatase in Neurospora crassa. mBio 12, e00871–21 (2021).

65. K. Lee, J. J. Loros, J. C. Dunlap, Interconnected Feedback Loops in the Neurospora Circadian System. Science 289, 107–110 (2000).

66. A. Padlom, et al., Level of constitutively expressed BMAL1 affects the robustness of circadian oscillations. Sci Rep 12, 19519 (2022).

67. Y. O. Abe, et al., Rhythmic transcription of Bmal1 stabilizes the circadian timekeeping system in mammals. Nat Commun 13, 4652 (2022).

68. B. Wang, A. N. Kettenbach, S. A. Gerber, J. J. Loros, J. C. Dunlap, Neurospora WC-1 Recruits SWI/SNF to Remodel frequency and Initiate a Circadian Cycle. PLoS Genet 10, e1004599 (2014).

69. Vogel, H.J. (1956) A Convenient Growth Medium for Neurospora crassa,. Microbial Genetics Bulletin 13, 42–47.

70. B. Wang, J. C. Dunlap, Domains required for the interaction of the central negative element FRQ with its transcriptional activator WCC within the core circadian clock of Neurospora. Journal of Biological Chemistry 299, 104850 (2023).

71. B. Wang, X. Zhou, S. A. Gerber, J. J. Loros, J. C. Dunlap, Cellular Calcium Levels Influenced by NCA-2 Impact Circadian Period Determination in Neurospora. mBio 12, e01493–21 (2021).

72. B. Wang, et al., A crucial role for dynamic expression of components encoding the negative arm of the circadian clock. Nat Commun 14, 3371 (2023).

73. K. Lee, J. C. Dunlap, J. J. Loros, Roles for WHITE COLLAR-1 in Circadian and General Photoperception in Neurospora crassa. Genetics 163, 103–114 (2003).

74. V. D. Gooch, et al., Fully Codon-Optimized luciferase Uncovers Novel Temperature Characteristics of the Neurospora Clock. Eukaryot Cell 7, 28–37 (2008).

75. H. V. Colot, et al., A high-throughput gene knockout procedure for Neurospora reveals functions for multiple transcription factors. Proc. Natl. Acad. Sci. U.S.A. 103, 10352–10357 (2006).

